# Effect of host-switching on the eco-evolutionary patterns of parasites

**DOI:** 10.1101/2021.11.27.470149

**Authors:** Elvira D’Bastiani, Débora Princepe, Flavia MD Marquitti, Walter A Boeger, Karla M Campião, Sabrina LB Araujo

**Author notes:** Corresponding author: Elvira D’Bastiani.

## Abstract

Increasing empirical evidence has revealed that host-switching are common in the history of parasites. Still, few have explored how the evolutionary histories of hosts might influence such switches and then the evolution of parasites. Here, we investigated how the intensity of host-switching, assumed to depend on the phylogenetic distance between hosts, affects the ecological and evolutionary patterns of parasite species. We developed an individual-based model where parasites can explore and colonise hosts under variable host-switching intensity and have evolution driven by mutation, genetic drift, and mating restriction. We hypothesised that our model can reproduce ecological and evolutionary patterns of empirical communities, characterised by turnover among host species and tree imbalance, respectively. We found an optimum range of host-switching intensity that can predict similar patterns as those observed in the empirical studies, validating our hypothesis. Our results showed that the turnover decreased as the host-switching intensity increased with low variation among the model replications. On the other hand, the tree imbalance had not a monotonic tendency but a wide variation. These results revealed that while the tree imbalance is a sensitive metric to stochastic events, the turnover may be a proxy for host switching. Furthermore, local empirical studies corresponded to higher host-switching intensity when compared to regional studies, highlighting that spatial scale is probably the crucial limitation of host-switching.

## Introduction

The dispersal of parasite individuals followed by colonisation of a new host lineage, known as host-switching, is a common event observed during the evolutionary trajectory of many parasite lineages (De Vienne et al. 2013). Initially, host-switching results in the increase of the host repertoire of a parasite (Braga et al. 2021). The colonisation of the new hosts can result in reproductive isolation, and consequently in speciation of parasite lineages, characterising the dynamics of the Oscillation Hypothesis (Nylin and Soren 2018). Empirical examples showing high levels of host-switching include symbiotic interactions ranging from host-parasite (Meinilä et al. 2004; Agosta et al. 2010; Müller et al. 2018, Fecchio et al. 2019; Boyd et al. 2022) and plant-insect systems to microbial pathogens (Woolhouse et al. 2005), brood parasitism (Habermannová et al. 2013; Dominguez et al. 2015), plant-feeding insects, and parasitic plants (Nylin et al. 2014). Consequently, understanding the factors influencing the success of host-switching and subsequent speciation events is critical for understanding parasites diversification.

A general framework that has been used to understand infectious disease, the Stockholm Paradigm, explores the evolutionary dynamics of host-parasite associations (Brooks et al. 2014; Brooks et al. 2019). This framework suggests that parasites perform host-switching by ecological fitting hypothesis (Agosta and Klemmens 2008; Agosta and Brooks 2020). Ecological fitting explains how the process whereby organisms colonise and persist in novel environments, use novel resources, or form novel associations with other species through a set of traits/capabilities they already possess (see Agosta and Klemmens 2008; Brooks et al. 2014; Brooks et al. 2019; Agosta and Brooks 2020). The expression of these unexplored capabilities is mediated by the opportunity of interaction (temporal and spatial) and determines the possibility of encounters between hosts and unfamiliar parasites. After the encounter, and if the interaction is compatible, it is followed by the resolution of subsequent conflicts that emerge from the basic dynamics of “living together”, which should result in co- accommodation (Brooks and McLennan 2002; Araujo et al. 2015).

Ecological and life-history traits also influence the chances of parasites dispersing from one host species to another. Characteristics of all organisms within the interaction system, such as niche similarity among host species, modes of transmission of parasites, dietary preferences of the vector (if there is one), and also ecosystemic characteristics as the host community composition and shared phylogenetic history are relevant factors that define the chances of host-switching (Bush et al. 2006; Jaramillo and Rivera-Parra 2018). Niche similarity among host species is one fundamental element constraining the incorporation of new host species by ecological fitting. This is because the capacity of a parasite species to use new resources is related to the phylogenetic conservatism of the resource provided by the host species. Phylogenetic distance between the original and new host species can represent an adequate proxy for the nature of the resource, which is tracked by the parasite lineage (Charleston and Robertson 2002; Agosta and Klemmens 2008; Engelstädter and Fortuna 2019). Consequently, the host phylogenetic conservatism can define the arena of possibilities for host-switching.

Several studies have indicated the ubiquity of host-switching in nature (see Cuthill and Charleston 2013; De Vienne et al. 2013; Engelstädter and Fortuna 2019; Fecchio et al. 2019; Hayward et al. 2021), but (or yet) few studies have explored to which extent the switches are constrained by inherited possibilities and limitations across hosts evolutionary histories. Among many potential factors determining host-switching, it seems that host phylogeny and geographic distributions are two major players (Sanaei et al. 2021). Moreover, the relation between host-switching and the opportunity for parasite dispersal, as well as their capacity to explore new hosts, is mostly unexplored (Brooks et al. 2019). Here, we aim to fill these unexplored gaps by proposing a novel approach to investigate how the intensity of host-switching affects the ecology and phylogenetic history of the parasites. For this, we assume that compatibility and the opportunity for interaction (spatial and temporal) may be expressed through the evolutionary histories of the hosts, and this can influence the host-switching events.

In this study, we propose a theoretical model based on parasite individuals that can switch among host species and speciate over time. Host-switching is mediated by phylogenetic conservatism; that is, the probability of parasites switching hosts decreases with increasing divergence in the evolutionary time of the hosts. The overall intensity of host-switching is a controlled parameter of the model. Under the absence of host-switching, the model is adjusted to parasites speciate due to limitation of host use, resulting in a pattern of cophylogeny and in paired specialised interaction (each parasite species interacting with one exclusive host species). We then investigate the eco-evolutionary patterns under different host-switching intensities, hypothesising that there is an optimum range of host-switching intensity that can result in the same eco-evolutionary patterns observed in the empirical studies. These patterns were characterised by species interaction turnover and tree imbalance, respectively. The model predictions were compared to nine empirical communities, validating our hypothesis.

## Material And Methods

### The model

We performed simulations of eco-evolutionary trajectories of parasites influenced by their host evolutionary history and host-switching events using an individual-based model (IBM). We assumed that the evolutionary history of the host can represent a proxy for the resources for parasite species (Agosta et al. 2010; Imrie et al. 2021), and also assumed that the probability of host-switching decreases as the phylogenetic distance between the species of host involved in the event (original and new host species) increases (Araujo et al. 2015; Engelstädter and Fortuna 2019). The model assumes that parasite evolution occurs at the same evolutionary time scale as the host, which increases possibilities for host-switching as host speciation occurs (Fig. 1a-c).

**FIGURE 1.**
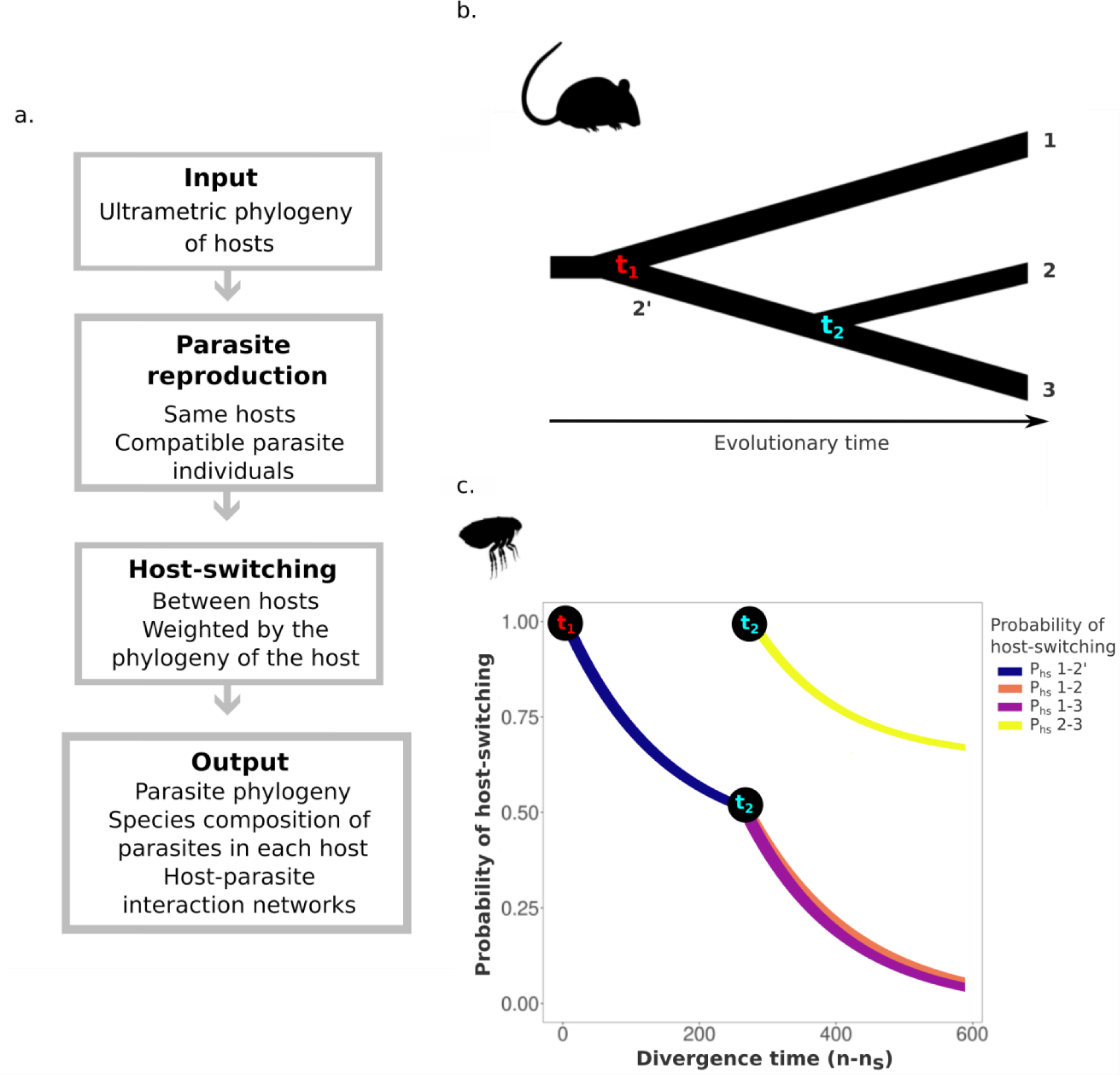
Schematic representation of the model. **a**. The general sequence of the model dynamics. **b**. Hypothetical host phylogeny. **c**. Probability of host-switching (P_hs_) over time. Each parasite individual can host-switch after the first speciation event (t_1_). One host is drawn for each parasite individual, and the probability of a successful host-switching depends on the divergent time between the two involved hosts. At t_1_ the first speciation event occurs, and the probability of host-switching is maximum. As time goes on, this probability decreases. At t2 another speciation event occurs, increasing the number of migration possibilities. At this time the two younger host species (2 and 3) have the maximum probability of switching hosts (P_hs_ 2-3), but the probability of host-switching between 1 and 2 or between 1 and 3 keeps decreasing. The colors highlight the 2’, 1, 2, and 3 host lineages presented in **b**.

Parasite individuals are explicitly described by biallelic sequences of infinite sites, a simplified form to represent their genomes and heritable trait. Individuals are monoic and engage in sexual reproduction, with non-overlapping generations, following the model proposed by Higgs and Derrida (1991) and Manzo and Peliti (1994). Population evolution is driven by mutation, genetic drift, and restriction to mating in the absence of natural selection. With a certain set of parameters, parasite speciation occurs. Each parasite individual is also characterised by the host species that it interacts. The host species are modelled as resources that impose a carrying capacity of *K* parasite individuals, analogous to islands in the Manzo-Pelit model (Manzo and Peliti 1994), but, in our model, the islands (hosts species in our case) emerge (as a new host species that speciate) according to a predetermined host diversification time (i.e. based on ultrametric empirical phylogenies - an ultrametric tree is a kind of additive tree in which the tips of the trees are all equidistant from the root of the tree). Thus, the overall carrying capacity increases by *K* individuals at each new host speciation. The model does not consider the selection pressure imposed by parasites on the evolution of the resource (host). Therefore, we are not modelling a process of reciprocal evolution, or co-evolution.

### Reproduction of parasites

Reproduction is sexual and occurs between parasite individuals that are in the same host and that have a minimum genetic similarity, q_min_, measured based on the Hamming distance between genomes. In each host species, at each generation, *K* offspring individuals replace the parental population, with no generation overlapping. We establish a maximum of *K* random trials with reposition to find one compatible partner. The offspring is generated by *locus* recombination of the parents and each *locus* has a probability of mutation (μ). We set *q_min_* = *0.5q_0_*, where *q_0_* is the expected mean similarity within one population in equilibrium: 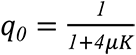. For a detailed demonstration of the above equation see SI1. The restriction *q_min_* = *0.5q_0_* is arbitrary and only assures that no parasite speciation occurs when using a unique host (i.e., avoids sympatric speciation in the context of Higgs and Derrida (1991)). Consequently, parasite speciation only happens when more than one host species is used.

### Temporal scaling

The empirical studies have evolutionary times in the order of millions of years, and to maintain this time scale in the model would demand a high computational cost. As proposed by Costa et al. (2019), in our approach we adopted a high value of mutation rate (μ=0.025) in order to decrease the number of iterations (time steps or generations) necessary for speciation to occur. Furthermore, we assumed that, due to the shorter life cycle of parasites, they have a faster speciation rate when compared to their hosts (Dowton and Austin 1995; Light and Hafner 2007). To satisfy these conditions, we rescaled the whole host phylogeny assuming that the smaller branch length consists of the minimal time for parasites to speciate due to isolation by host use (see the demonstration in SM1):

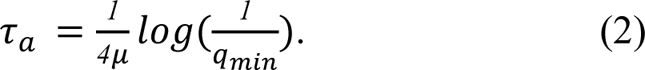

The minimal time for speciation decreases with *q_min_*. Therefore, making the reproduction more restricted (i.e., increasing *q_min_*) facilitates the formation of parasite species in a shorter time. The simulation starts with a clonal parasite population using a unique host species. Also, the first host speciation occurred only after *τ_a_* generations for the parasite populations to accumulate genetic diversity before the first splitting event.

### Host-switching events

After the first host speciation, parasite individuals in a host species may switch to another host. For each parasite individual, we randomly selected a host species, including the one in use. If the selected host species is not the original host (donor), we follow a probability function for the host-switching event. This probability of host-switching events (*P_hs_*) decreases over time, representing the product of opportunity for contact and compatibility of the interaction of parasites associated with the evolutionary history of hosts (Fig. 1c). Then, we are assuming that compatibility, the opportunity of interaction are expressed through the evolutionary history of hosts. The probability of a parasite individual successfully migrates (host-switching) from one host to another host species, in a given generation *n*, is defined as:

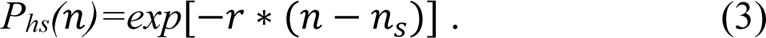

where *r* is a positive parameter that controls the decay of the host-switching probability, and *n_s_* is the generation that the common host ancestor had speciated (then, n-*n_s_* is how long the two host species had diverged). If *r = 0,* these probabilities are equal to 1 regardless of the host divergence time, meaning that there is no restriction to host-switching. As a consequence, parasite gene flow is continuous and speciation does not occur. At the other extreme, for sufficiently large *r* values (P*_hs_*∼0), host-switching is absent and cospeciation between hosts and parasites is expected. For intermediary *r* value, some parasite individuals can eventually switch hosts (Fig. 2 and Fig. S1). This will increase the host repertoire of the parasitic species, and also enable speciation by isolation (by host use), similar to the speciation by founder’s effect (Mayr 1999; Gavrilets and Hastings 1996). The effect of the overall host-switching in a community does not depend only on *r*, but also on the particularities of each host phylogeny that is used as input for the calculation of the host-switching probability. Therefore, to better interpret the effect of parameter *r* on the trajectories and compare the results between the communities, we do not present our results in terms of *r,* but how much it changes the overall host-switching events. To obtain this overall metric, we calculated the mean percentage of parasite individuals that switch hosts over the entire simulation and we call it ***host-switching intensity*** (Fig. 2).

**FIGURE 2.**
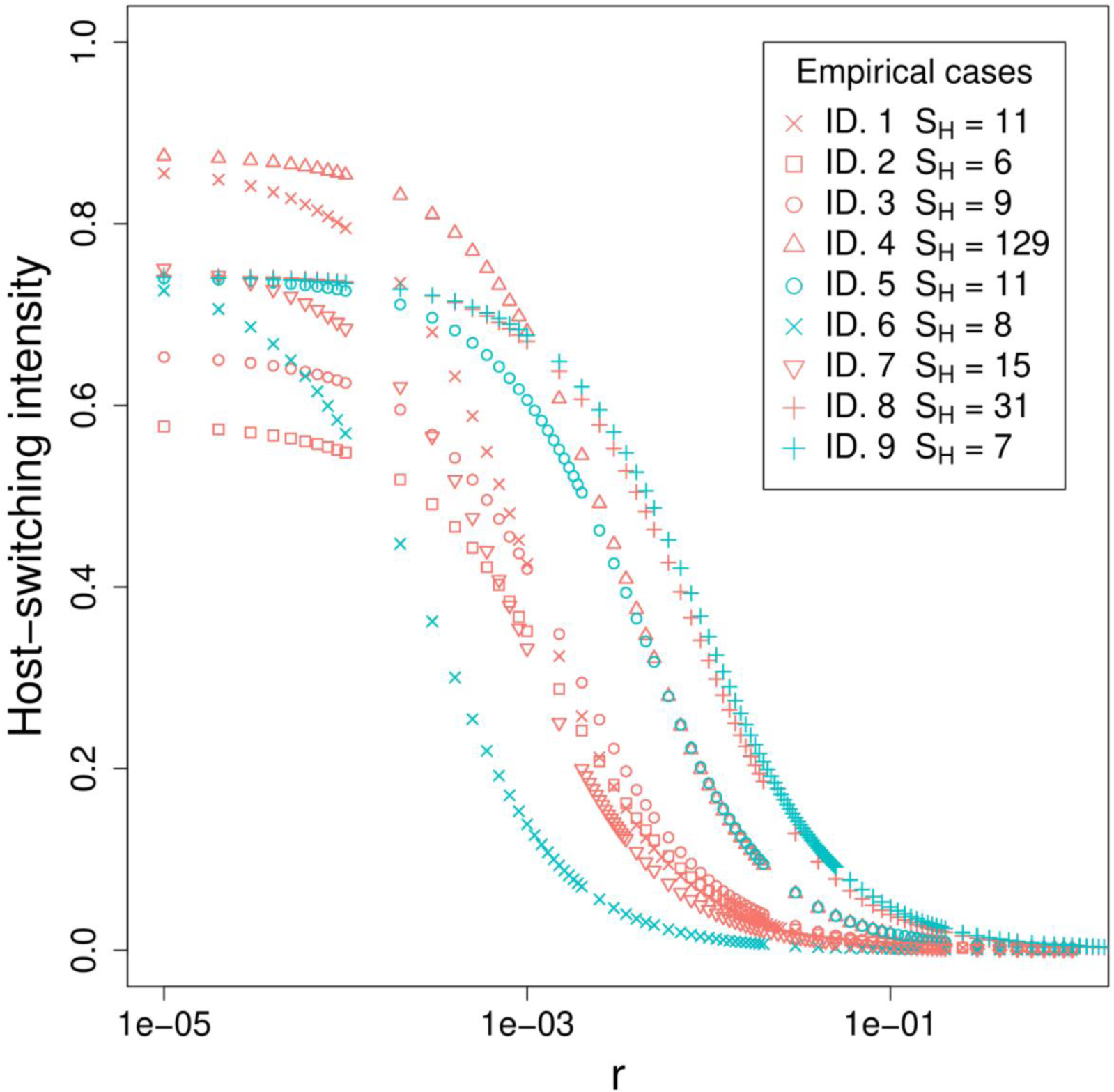
Relation between *r* (a parameter that defines the host-switching decay, Eq. 3) and the intensity of host-switching for each simulated community. Each ID represents empirical studies and S_H_ represents the host richness. ID. 1 - Birds and feather mites. ID. 2 - Mammals and lice. ID. 3 - Wildlife and ectoparasites. ID. 4 and 5 - Rodents and fleas. ID. 6 - Fish and Monogeneans (Gyrodactylidae). ID. 7 - Frogs and monogeneans (Polystomatidae). ID. 8 -Frogs and lungworms (*Rhabdias* spp.). ID. 9 - Frogs and gut worms (*Oswaldocruzia* spp.). Regional spatial scale studies are represented by salmon colour and local spatial scale studies by blue.

### Parameters of the model

For the results presented here we fixed the population size per host (*K* = 250) and mutation probability per locus (µ = 0.025). With these parameters, we can observe species formation with reasonable computational time. Since the empirical studies varied in the number of host species and branch size, the total number of iterations also varied (Table S1). The parameter *r* varied (*0<r<1*) for each empirical study. A total of 50 replicates were performed for each parameter combination. We have also analysed the model predictions under other values of population size (*K* = {50, 500,1000}) and mutation rate (μ = 0.001) (Table S1). Our qualitative conclusions did not change under these parameter variations (Fig. S2-S4).

### Validation with empirical data

The development of a new method to assess the host-switching intensity allowed us to compare the results of our simulations with empirical data from different groups of parasites and their respective hosts. This method uses information on the evolutionary history of the host species as a proxy for resource similarity. We used nine studies from empirical studies of parasite-host associations (Table 1) for comparative purposes. The selection criteria was that, in addition to information on species interaction, these empirical studies essentially needed to have phylogenies for hosts and parasites (see the details in Fig. S5-S13). We separated these empirical studies according to the spatial scale (Table 1). Spatial scale refers to the spatial extent of ecological processes and the spatial interpretation of the data. In this study, we assumed that studies in the local spatial scale are essentially in a geographic radius less than or equal to 35km, while on a regional scale they were collected essentially in a geographic radius greater than 35km, in the original article respectively.

**TABLE 1.** Description of the host sample size and parasite richness for each empirical study, of which host phylogenies were used as model parameters and host-parasite association to validate the simulations of the model. Legend: ID = Empirical study.

| ID | Host group | Host richness | Parasites group | Parasite richness | Spatial scale | Reference |
| --- | --- | --- | --- | --- | --- | --- |
| 1 | Bird | 11 | Feather mites<br>(*Trouessartia* spp.) | 11 | regional | Donã et al. 2017 |
| 2 | Mammals | 6 | Lice<br>(*Pediculus* spp. and<br>*Pthirus* spp.) | 7 | regional | Reed et al. 2007 |
| 3 | Wildlife | 9 | Arthropods* | 8 | regional | Becker et al. 2018 |
| 4 | Rodents | 129 | Fleas* | 202 | regional | Krasnov et al.<br>2016 |
| 5 | Rodents | 11 | Fleas* | 19 | local | Krasnov et al.<br>2016 |
| 6 | Fish | 7 | Monogeneans<br>(Gyrodactylidae) | 16 | local | Patella et al. 2017 |
| 7 | Frogs | 15 | Monogeneans | 13 | regional | Badets et al. 2011 |
| 8 | Frogs | 31 | Lungworms<br>(*Rhabdias* spp.) | 18 | regional | Müller et al. 2018 |
| 9 | Frogs | 7 | Gut worms<br>(*Oswaldocruzia* spp.) | 5 | local | Willkens et al.<br>2021 |

Note: \*Include different parasite groups

### Characterization of the ecological and evolutionary patterns of parasites

We compared both the structure of turnover of parasite species (ecological pattern) and the imbalance of parasite phylogenies (evolutionary pattern) in the empirical studies with those resulting from the simulations. To characterise the composition of parasite species we used the metric that gives information about the beta diversity of multiple-site dissimilarities (*β_SOR_* - Baselga 2010; 2013a, b). The beta diversity may reflect two different phenomena: turnover (*β_SIM_*) and nestedness (*β_NES_*) (Baselga et al. 2007; Baselga 2010; 2013a, b). Here, we choose only to work with the Simpson-based multiple-site dissimilarity, that is turnover (*β_SIM_*), since it is non-dependent on species richness (Baselga et al. 2007; Baselga 2010). This refers to the replacement of some species by others as a consequence of environmental sorting or spatial and historical constraints. In our case, we compared the variation in parasite species composition between host species. The Simpson-based multiple-site dissimilarity is then:

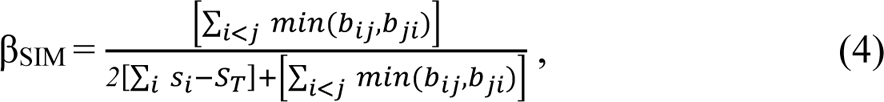

where *S_i_* is the total number of species in site *i*, *S_T_* is the total number of species in all sites (hosts in our case) and min(*b_ij_,b_ji_*) is the minimal number of species exclusive to sites *i* and *j* in pairwise comparison (Baselga 2010).

To characterise the structure of the phylogenetic trees we used the metric that gives information about the tree imbalance. Tree imbalance is one of the most common phylogenetic structural patterns and measures asymmetries between the numbers of species on each side of the tree’s branches (Marquitti et al. 2020). Tree imbalance is widely measured using the Sackin index (*I*) (Sackin 1972; Blum and François 2005; Frost and Volz 2013; Dearlove and Frost 2015). The *I* has a dependence on the number of leaves, making it unsuitable for comparing trees with different numbers of species. To make this comparison possible, we use the normalised Sackin index (*I_n_*) given by:

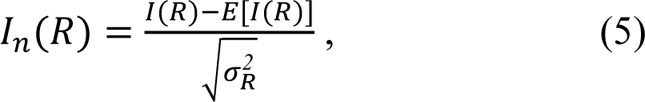

where tree imbalance is the *I(R)*, and *E(I_n_)* and 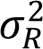 the expected and variance of trees generated by the Yule model which have the same number of leaves (species) as the observed tree (Cardona et al. 2013; Marquitti et al. 2020). Although I_n_(R) would be close to zero for trees generated with the Yule model, independent of the species richness R, different modes of speciation may introduce important deviations from the behaviour of the Yule model (Marquitti et al. 2020).

As each empirical study represents particular ecological and evolutionary processes, we analysed whether there was an optimal range of host-switching intensity in our simulated cases that retrieves information about turnover (*β_SIM_*) and normalised Sackin index (*I_n_*) of each study. We considered that simulations that reproduced both the *β_SIM_* and the *I_n_* metrics simultaneously (within a ±*5%* confidence interval) were the best fit to the empirical examples. Then we compared the best fitting of host-switching intensity among the empirical studies to understand how it varied for different evolutionary histories. Although species extinctions occur in the model, this aspect was not included in the analyses since we do not have information about extinctions in the empirical studies. These analyses were performed using ‘ape’ (Paradis and Schliep 2019), ‘betapart’ (Baselga et al. 2018) ‘picante’ (Kembel et al. 2010), ‘phytools’ (Revell 2012), and ‘vegan’ (Oksanen et al. 2013) R packages. See the details in SI3.

### Statistical analysis

To test whether the spatial scale modulates the best fitting host-switching intensities, a linear mixed-effects model (LMM) was performed using the *lmer* function from the ‘lme4’ package (Bates et al. 2015). We assumed the host-switching intensity as the response variable, the spatial scale as a fixed variable, and empirical studies were treated as random variables (intensity∼ scale+(1|study)). After performing the LMM analysis, an analysis of variance (ANOVA) was used to determine significant differences (p-value ≤ 0.01) using the Anova function in the ‘car’ package (Fox and Weisberg 2019). All statistical analyses were performed in R v.4.0.0 (R Core Team 2020) and Rstudio v.1.3.959 (RStudio Team 2020).

## Results

The turnover and normalised Sackin index of parasites varied according to the mean percentage of parasite individuals that switch hosts during the entire history of the host community (the host-switching intensity). To illustrate the turnover and normalised Sackin index according to the host-switching intensity, we present an example of a model application with fleas associated with rodents (ID. 5, Fig. 3a-c). As expected, turnover decreases as host-switching intensity increases (Fig. 3a and Fig. S14). This occurs because the increase of host-switching promotes the interaction of different host species with the same parasite species.

**FIGURE 3.**
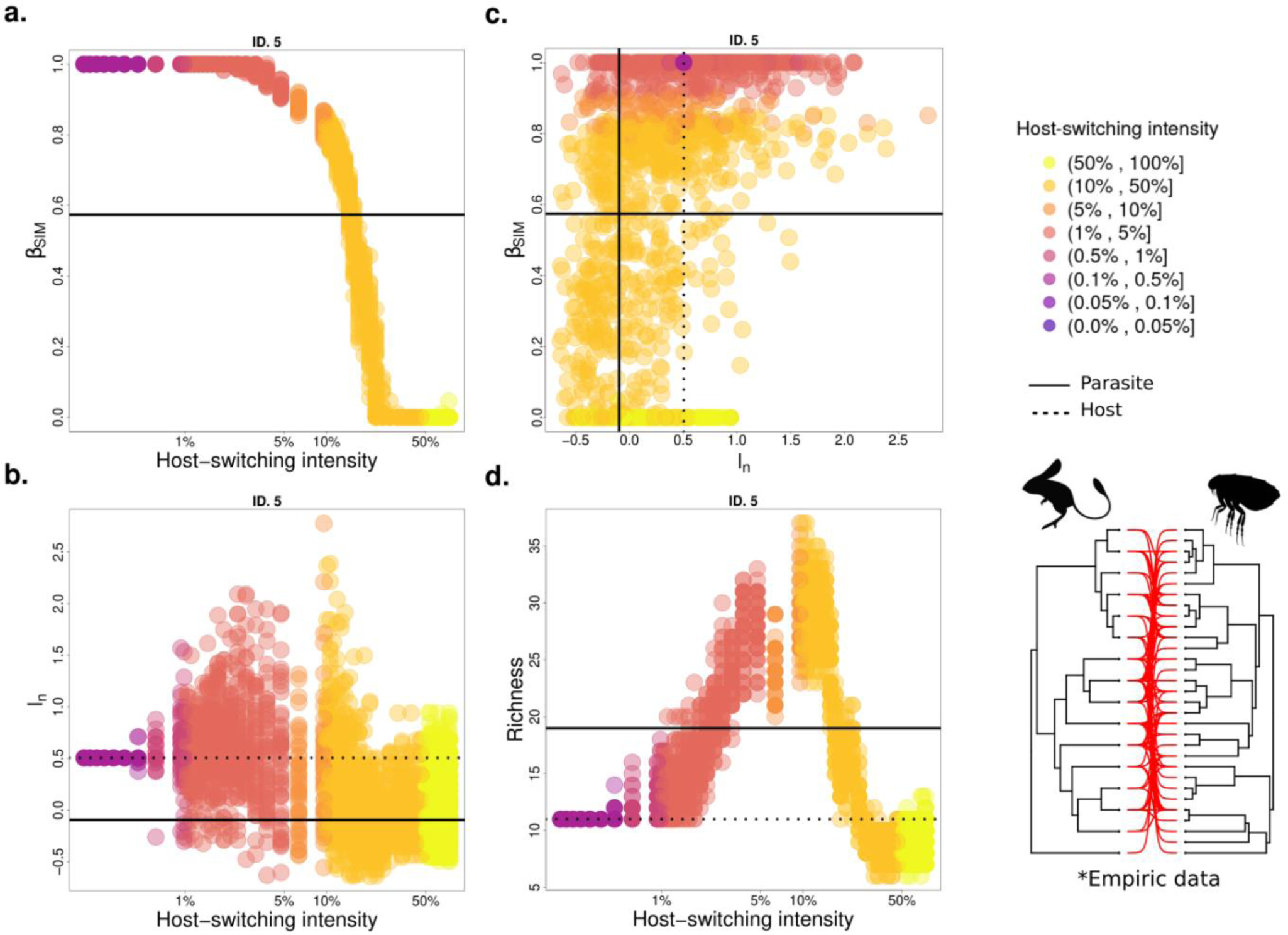
Influence of host-switching events on the eco-evolutionary patterns of simulated parasites for fleas associated with rodents (see Table 1 for details). Here we demonstrated the relationship between: **a**. Host-switching intensity and turnover of parasite species (β_SIM_) between host species; **b**. Host-switching intensity and parasite normalised Sackin index (I_n_); **c**. Relationship between β_SIM_, I_n_, and the host-switching intensity. **d**. Host-switching intensity and parasite richness. The lines refer to empirical information of parasite (continuous) and host (dotted). The colored dots are redundant with the x-axis scale of graphs (a) and (b) but intend to guide the interpretation of (c). A total of 50 replicates were performed with 250 individuals for each configuration of the parameters of host-switching intensity.

Additionally, for each value of host-switching intensity, there is a small variation in the turnover (Fig. 3a and Fig. S14). The only exception was ID. 4, which resulted in a wide variation in turnover under high host-switching intensity (Fig. S14).

As imposed by the model, the parasite richness ends the same as hosts in the absence of host-switching (Fig. S16). But, for intermediary values of host-switching, parasites can colonise the new host and then speciate, resulting in an overwhelming increase in parasite speciation (see the dynamics in the movie available in S17, Fig. S18, and Fig. 3d).

When host-switching intensity is low (below 1%), the normalised Sackin index (I_n_) for the simulated parasite phylogenies results in the exactly same value as the one obtained from the empirical phylogeny of the host (note the dashed line in Fig. 3b and also Fig. S15). This is because the low host-switching intensity does not allow the establishment of the parasite in a new host and, as a consequence, the simulated parasite phylogenies have the same normalised Sackin index of the empirical host phylogeny. Colonisation followed by speciation is more likely to occur under a higher host-switching intensity, in which the normalised Sackin index varies over simulations even when they are under the same host-switching intensity (Fig. 3b and Fig. S15). The wide variation in the normalised Sackin index for a given host-switching intensity reveals that stochastic host-switching events, even if host-switching is more likely to occur between closely related species, can change the structure of the resulting phylogenetic tree. Despite not having a monotonic tendency, the normalised Sackin index tends towards zero (balanced tree) as host-switching intensity goes to one, regardless of the community (Fig. S15), resembling a neutral speciation scenario Yule model (Yule 1924; Aldous 2001).

For all empirical studies analysed, there is a range of host-switching intensity that simultaneously reproduces the observed turnover and the parasite normalised Sackin index (Fig. 4). As mentioned, both metrics are sensitive to host-switching intensity but each one varies independently of the other (see in Fig. 4). Generally, the turnover and the parasite normalised Sackin index obtained under high host-switching intensity (greater than 50%) are far from the empirical pattern (see Fig. 4, the yellow dots rarely approach the intersection of the solid lines).

**FIGURE 4.**
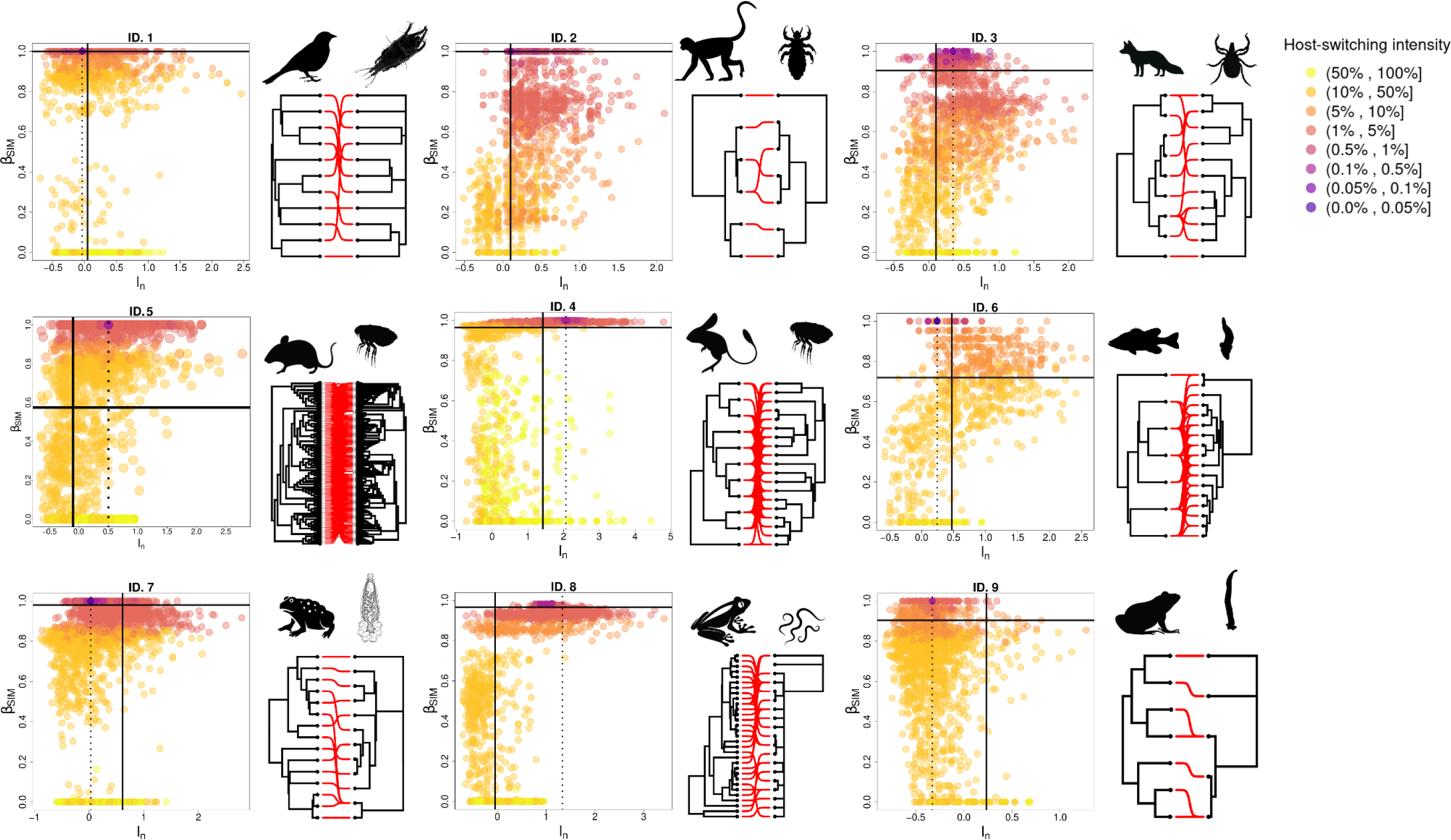
The relationship between variation in the turnover of parasite species, normalised Sackin index of parasite species, and host-switching intensity for nine empirical studies: the parasites’ turnover, measured by the metric turnover(β_SIM_) on the y-axis and the normalised Sackin index (I_n_) on the x-axis. Each ID represents an empirical case. The lines refer to empirical information of parasite (continuous) and host (dotted). Colour scales represent each percentage interval of host-switching intensity. A total of 50 runs were performed with 250 individuals of carrying capacity, for each configuration of the parameters of host-switching intensity.

The simulated host-switching intensity that simultaneously fit parasite turnover and normalized Sackin index recovered a range of 0.06% to 22.07% of host-switching intensity through the analysed empirical studies. Within this range, the associations between mammals and lice presented the lowest host-switching intensity (case ID. 2 with 0.07% - 1.13%), followed by that involving wildlife and arthropod parasites (case ID. 3 with 0.43% - 2.69%), frogs and monogeneans (case ID. 7 with 0.22% - 3.71%), frogs and lungworms (case ID. 8 with 1.99% - 4.94%), frogs and gut worms (case ID. 9 with 5.29% - 9.35%), birds and feather mites (case ID. 1 with 0.06% - 8.17%), fish and monogeneans (case ID. 6 with 8.26% - 11.64%), - the highest intensities of host-switching were observed between rodents and fleas (case ID. 5 with 14.45% - 16.87% and case ID. 4 with 0.43% - 22.07%). We also observed that the host-switching events are more frequent in studies conducted in a local scale (blue colour in Fig. 5) than in regional scales (salmon colour in Fig. 3) (LMM: relationship of host-switching intensity on spatial scale: beta= 0.08, SE= 0.01, df = 6.92, t= 5.25, p = 0.001, ANOVA: F = 27.56, p = 0.001, Fig. 5).

**FIGURE 5.**
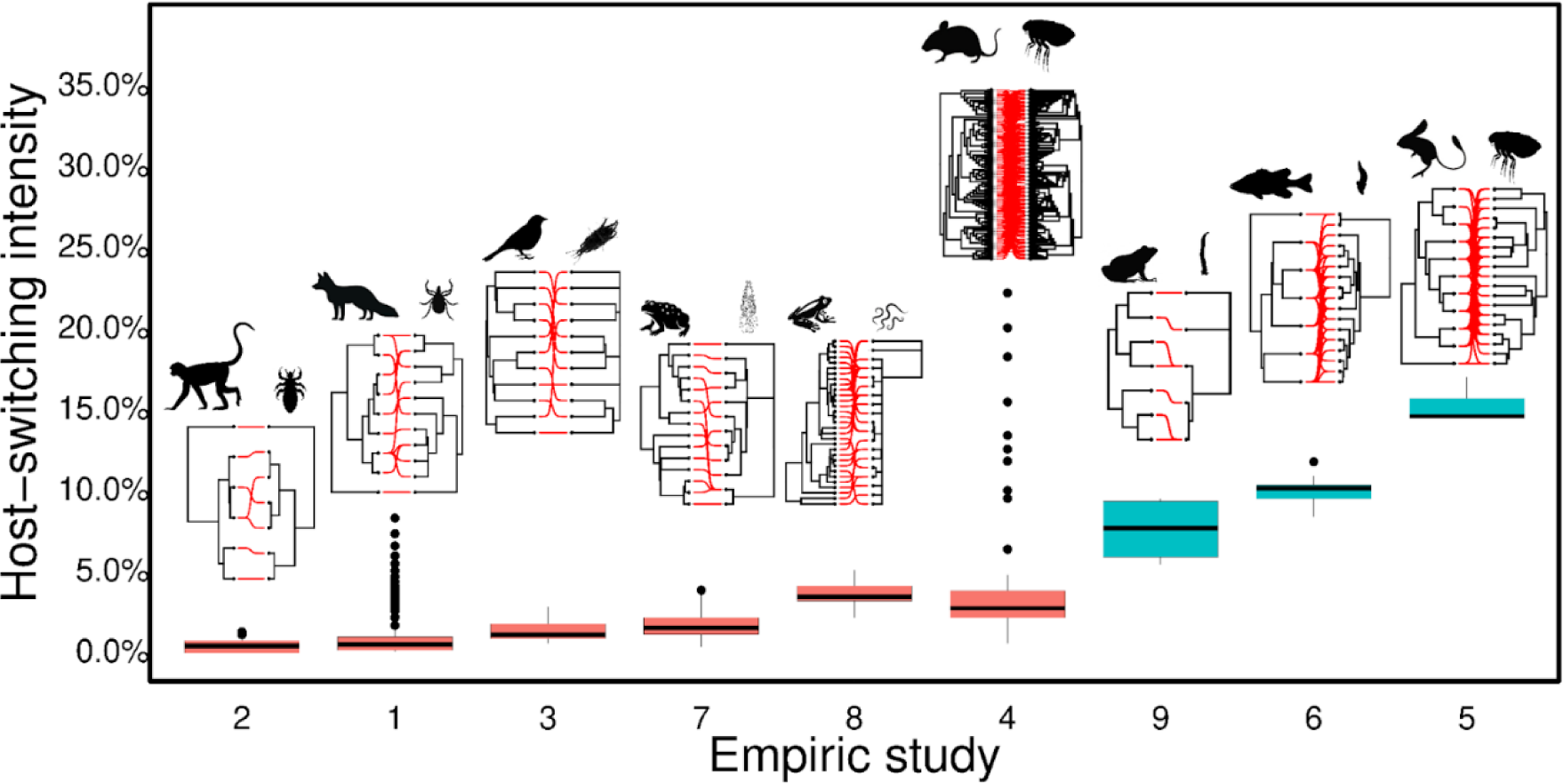
Calculated host-switching intensity among empirical studies. The boxplots show the distributions of simulated host switching intensities for each empirical study. Regional spatial scale studies are represented by salmon, and local scale studies are in blue. The number in axis x represents the empirical studies: ID. 1 - Birds and feather mites. ID. 2 - Mammals and lice. ID. 3 - Wildlife and ectoparasites. ID. 4 and 5 - Rodents and fleas. ID. 6 - Fish and Monogeneans (Gyrodactylidae). ID. 7 - Frogs and monogeneans (Polystomatidae). ID. 8 - Frogs and lungworms (*Rhabdias* spp.). ID. 9 - Frogs and gut worms (*Oswaldocruzia* spp.).

## Discussion

In this study, we developed a novel methodological framework to understand how the intensity of host-switching shapes some aspects of ecological and evolutionary patterns of parasites, here characterised by species interaction turnover and tree imbalance, respectively. Our three main results are 1) We found an optimum range of host-switching intensity that can predict similar patterns as those observed in the empirical studies, which validates our model; 2) The model showed that the increase of host-switching intensity promoted an increase in turnover, but the tree imbalance did not follow any monotonic tendency. Moreover, for a specific host-switching intensity, we observed a small variation in the turnover and a wide variation in the tree imbalance; 3) The predicted values of host-switching intensity varied among the empirical studies and those at a local spatial scale resulted in values higher than the ones at the regional scale.

The fact that our model rebuilt the eco-evolutionary patterns of all empirical studies supports the idea that host-switching mediated by host evolutionary proximity is a good predictor of parasite associations. According to the framework of the Stockholm Paradigm (Brooks et al. 2019), parasites can colonise new host species due to pre-existing compatibility, which is expressed when there is an opportunity for contact. Compatibility emerges greatly from the ancestral capacity in which both hosts and parasites must be physiologically compatible to establish a long-term association (Brooks and McLennan 2002; Kolbe et al. 2004; Brooks et al. 2019). Hence, for a given lineage of the parasite, the closer (phylogenetically) the original and the new host species, the greater the possibility that the adequate resource is conserved or is at least similar. In fact, phylogenetic proximity has been widely recognized as a potential criterion to anticipate new associations (Streicker et al. 2010; Damas et al. 2020; Filion et al. 2022).

Another element of the Stockholm Paradigm (Brooks et al. 2019) that we observed over the temporal dynamics of our model is the Oscillation Hypothesis (Janz and Nylin 2008): parasites first increase their host repertoire (generalise) and then speciate (specialise). In our model, at each time step, a parasite individual can switch hosts, promoting the increase of host repertoire for the parasite species. However, as we assume that the probability of host-switching decreases as hosts diverge, given time, the probability of individuals from the same parasite species maintaining the gene flow between those host species decreases, and parasite speciation is likely to occur (see the dynamics in the movie available in S17 and S18). Consequently, in our model, host-switches promote host repertoire oscillation, as hypothesised by Janz and Nylin (2008), and favour parasites to speciate at a greater rate than their host, which is empirically evidenced (Poulin and Morand 2000). Although the mean argument behind the difference in speciation rate between host and parasite is the parasite’s shorter life cycle, we support the idea that frequent host exploitation is another important mechanism to parasite diversification (Hay et al. 2020, Boeger et al. 2022).

The tree imbalance did not have a monotonic tendency and showed a wide variation for a given host-switching intensity. This reveals that stochastic events can change the evolutionary trajectory of parasites. Although our model assumed that host-switching most likely occurs between closely related species, eventually, a parasite can switch to a phylogenetically distant host, changing the diversification history completely. This distant host-switching was observed in most of the empirical studies presented here, where the parasites were able to colonise hosts from different genera (ID 2 and 6), families (IDs 1, 4, 5, 7, 8, and 9), and even order (ID 3). For example, in study ID8 the *Rhabdias* lung-worm anuran parasites occurred mostly in Bufonidae hosts and only one species in the Hylidae host (Müller et al. 2018). Species extinction is another class of stochastic event present in our model that could contribute to the varied outputs on parasite evolution. As we use data only of extant species, when a species goes extinct all its history is lost, also impacting the imbalance of the tree (Costa et al. 2019, Marquitti 2020).

Unlike the tree imbalance, our results showed that the turnover has a monotonic tendency: it decreased as the host-switching intensity increased. This pattern was expected since the model imposes that as host-switching intensity increases, the limitation to use a new host decrease (Fig. 2). Moreover, we did not observe a wide variation in the turnover for a given host-switching intensity over the model replications. This reveals that those stochastics events mentioned before cannot produce significant changes in the turnover. It probably occurs because when host-switching occurs, it produces a decrease in the turnover no matter what parasite species switched to what host species. In other words, the identities of the species are not relevant since turnover emerges not from a given species characteristic, but from the similarities between species, or even spatial and temporal amplitudes (Fallon et al. 2004, Baselga et al. 2007, Baselga, et al. 2022). This reinforces the idea that species turnover is a robust metric to compare species assemblages (Baselga, et al. 2022) and may also be a good proxy for host-switching intensity.

The host-switching intensity varied across empirical studies and we observed that it is higher in empirical studies of local spatial scale than regional spatial scale. This evidence shows that the amplitude of the spatial scale is a fundamental factor in determining the extent of host-switching. The opportunity for interaction is larger in host empirical studies at a local scale, as this reduces the likelihood that barriers exist, hampering the encounter of potential actors. This is evident when comparing rodent and flea associations at regional (ID. 4) and local spatial scales (ID. 5). Similarly, since the association of *Rhabdias* spp. and frogs (ID. 8) are defined geographically (and not by host taxa) it was assumed that host-switching by ecological fitting was evolutionarily more important than association with particular host taxa (Kuzmin et al. 2014; Müller et al. 2018). Different intensities of host-switching observed in our results may also be influenced by biological variations of the species that make up the empirical studies analysed. For instance, these studies include a great diversity of organisms (fleas, lice, feather mites, helminths, platyhelminthes), with profound differences in their biological characteristics. Expanding analyses to a broader sample of empirical studies, including variations in the type of parasitism (e.g., mono *vs.* heteroxenic, ecto *vs.* endoparasite) and host attributes can provide important insights into key features related to the process of incorporation of new hosts.

In nature, host-parasite systems are more complex than those modelled here. Although the model can reconstruct eco-evolutionary patterns of empirical studies, our model has some limitations. For example, the carrying capacity of all host species is the same and the host’s body size, their abundances, and spatial distribution were not explicitly considered. The selective pressure is not explicitly modelled, contrary to what we observe in nature (Krasnov et al. 2005, Krasnov et al. 2021). Furthermore, all parasite individuals and species are equivalent, and may compete for the same resources. Except for resource competition, our model didn’t consider intra and interspecific interactions among parasites. Finally, the phylogenies are still scarce, especially for parasite species, which limited the number of tests with the model. Phylogenetic data on parasites is extremely important to clarify the role of host-switching in the ecological and evolutionary patterns of parasite lineages. Still, we recovered compatible eco-evolutionary patterns for modelled parasites and their respective hosts. Our model has important implications for predicting host switching, especially in scenarios of anthropogenic change. With anthropogenic changes constantly modifying natural environments and altering the geographic distribution of parasites, many species that were once restricted to specific areas are now expanding their range into new geographic locations and changing the composition of communities (see Brooks et al. 2014). As we showed, parasites can follow different evolutionary paths, and eventually can switch to non-related hosts, ultimately, determining the migration of a parasite to other species (and clades), in some cases, including humans. To conclude, we show that a model in which host-switching mediated by evolutionary proximity between hosts is a predictor for parasitic associations over evolutionary time, as well as for the origins of parasite diversity. We see this as an important step in our understanding of parasite diversification processes.

## Supplementary material

Data SI

## Data Availability

The model, phylogenies, and interactions of all analysed studies are available at https://github.com/elviradbastiani/host_switching_model.

## Acknowledgements

We thank the researchers who collected and reported the field data, our institutions, and the many colleagues who helped us in different ways during this project for their comments and suggestions, especially to Professor Dr. Marcus Aguiar. EDB is grateful for the Ph.D. degree scholarship provided by Capes (Coordenação de Aperfeiçoamento de Pessoal de Nível Superior). The authors acknowledge the computational support from Professor Carlos M. de Carvalho at LFTC-DFis-UFPR.

## Financial Support

EDB was supported by Brazilian Coordination for the Improvement of Higher Education Personnel (CAPES). DP was supported by the São Paulo Research Foundation (FAPESP), grants #2018/11187-8, #2019/24449-3, and #2016/01343-7 (ICTP-SAIFR). SBLA was supported by Conselho Nacional de Desenvolvimento Científico e Tecnológico (CNPq: #11284/2021-3).

## Authors Contribution

Conceived and designed the experiments: EDB, SBLA and DP

Performed the experiments and analysed the data: EDB, SBLA and DP

Wrote the paper: EDB, SLB, KMC, DP, WB, FMDM

Other contributions: EDB, DP, FMDM, WB, KMC and SLBA

## Conflict of Interest

The authors declare that they have no conflict of interest.

## Notes

https://github.com/elviradbastiani/host_switching_model

### Competing Interest Statement

The authors have declared no competing interest.

### Summary of Updates

Associate Editor Comments to the Author: The manuscript has been evaluated by two referees, who agree that this study on modeling host-parasite cophylogenetic patterns addresses relevant questions for the journal Evolution. The referees however raise serious concerns. I have to agree with referees that it is unclear what hypotheses are tested and what conclusions can be inferred from the work. For example, the conclusion L14 in the abstract just seems to be a tautology; furthermore, that - empirical patterns can be reproduced from the model- does not say much. Is the model of speciation tested? Is the influence of host switching on phylogenetic patterns tested? Are host-switching rates estimated? Is the influence of ecto versus endo parasites tested? I have to agree that neither goal seems to be achieved convincingly. Furthermore, I also agree that too many simplifying assumptions are made without justification or discussion. I do not understand the choice of the speciation model considered (which is only explained in supplemental text while it is essential for understanding the study): the authors assume a kind of allopatric speciation scenario (ie, reproductive isolation arising as a by-product of genetic divergence), while the whole study deals with host switching and therefore new host adaptation. Reproductive isolation arising as a byproduct of host adaptation would seem more appropriate (eg 10.1016/j.tree.2010.03.006). I also have to agree that it is unclear why two particular measures (beta diversity and tree imbalance) have been chosen to assess the fit of patterns between data and modeling and what they mean. More specific comments: -L41-42: I would not call this a theoretical framework or a paradigm, it just seems quite obvious from the evolutionary theory -L71-72, L80, L287: there are several studies in plant-fungus associations, eg https://doi.org/10.1111/j.1420-9101.2009.01878.x; https://doi.org/10.1073/pnas.0607968104 -L160: reproductive isolation following host switching has been specifically modelled and is not similar to speciation by founder s effect but rather to immigrant s inviability 10.1016/j.tree.2010.03.006 (ie, the barrier is intrinsic and not extrinsic); the choice of an allopatric speciation model in a study dealing with host switching seems weird.

