## SUPPLEMENTARY MATERIAL for "Effect of host-switching on the eco-evolutionary patterns of parasites"

### Data SI

METHODS

*Additional information for the model*

*Derrida-Higgs model*

The model introduced by Derrida and Higgs (1991) considers a sympatric population of *K* haploid individuals (population carrying capacity) whose genomes are represented by binary strings of size *B*, $\{S_1^\alpha, S_2^\alpha \dots S_B^\alpha\}$ where $S_i^\alpha$, can assume the values ±1. Each locus of the genome is dumbed as a gene and the values +1 and -1 the corresponding alleles. The number of individuals at each generation is kept constant and the population is characterised by a $K \times K$ matrix $q$ measuring the degree of genetic similarity between pairs of individuals:

$$q_{\alpha\beta} = \frac{1}{B}\sum_{i=1}^{B} S_i^\alpha S_i^\beta. \quad (1)$$

If the genomes of $\alpha$ and $\beta$ are identical $q_{\alpha\beta}$= 1 whereas two genomes with random entries will have $q_{\alpha\beta}$ close to zero. Each generation is constructed from the previous one as follows: a first parent $P_1$ is chosen at random. The second parent $P_2$ has to be genetically compatible with the first, i.e., their degree of similarity has to satisfy $q_{P_1P_2} \geq q_{min}$. Individuals $P_2$ are then randomly selected until this condition is met with $K$ trials. If no such individual is found, $P_1$ is discarded and a new first parent is selected. The offspring inherits, gene by gene, the allele of either parent with equal probability (sexual reproduction). The process is repeated until $K$ offspring have been generated. Individuals are also subjected to a mutation rate $\mu$ per gene, which is typically small.

*Dynamics*

To understand how the similarity matrix changes through generations, consider first an asexual population where each individual α has a single parent $P_1$ in the previous generation. The allele $S_i^{\alpha}$ will be equal to $S_i^{P(\alpha)}$ with probability $\frac{1}{2}(1+e^{-2\mu}) \approx 1-\mu$ and -$S_i^{P(\alpha)}$ with probability $\frac{1}{2}(1-e^{-2\mu}) \approx \mu$, so that the expected value is $E(S_i^{\alpha}) = e^{-2\mu} S_i^{P(\alpha)}$. For independent genes, the expected value of the similarity between $\alpha$ and $\beta$ is, therefore, $E(q_{\alpha,\beta}) = e^{-4\mu} q_{P(\alpha),P(\beta)}$. In sexual populations $\alpha$ and $\beta$ have two parents each, $P_1^{\alpha}$, $P_2^{\alpha}$ and $P_1^{\beta}$, $P_2^{\beta}$, and since each inherits (on average) half the alleles from each parent, we obtain

$$E(S_i^{\alpha}) = \frac{1}{2}(S_i^{P_1(\alpha)}\frac{1+e^{-2\mu}}{2} - S_i^{P_1(\alpha)}\frac{1+e^{-2\mu}}{2}) + \frac{1}{2}(S_i^{P_2(\alpha)}\frac{1+e^{-2\mu}}{2} - S_i^{P_2(\alpha)}\frac{1+e^{-2\mu}}{2})$$

$$= \frac{e^{-2\mu}}{2}(S_i^{P_1(\alpha)} + S_i^{P_2(\alpha)}). \quad (2)$$

It follows that, on average, the similarity between $\alpha$ and $\beta$ is

$$q_{\alpha\beta} = \frac{e^{-4\mu}}{4}(q_{P_1(\alpha),P_1(\beta)} + q_{P_2(\alpha),P_1(\beta)} + q_{P_1(\alpha),P_2(\beta)} + q_{P_2(\alpha),P_2(\beta)}), (3)$$

with $q_{\alpha\alpha} \equiv 1$. In the limit of infinitely many genes, $B \rightarrow \infty$, this expression becomes exact and the entire dynamics can be obtained by simply updating the similarity matrix.

If there is no restriction on mating, $q_{min} = 0$, we can demonstrate that the overlaps $q_{\alpha\beta}$ converge to a stationary distribution (Derrida & Higgs, 1991), as follows. At generation *t*, the probability that $\alpha$ and $\beta$ have one parent in common, $P_1^\alpha = P_1^\beta, P_1^\alpha = P_2^\beta, P_2^\alpha = P_1^\beta$ or $P_2^\alpha = P_2^\beta$, is *4/K*. In this case, the average similarity $< q_{\alpha\beta} >$ between $\alpha$ and $\beta$ is given by $< q_{\alpha\beta} >= \frac{e^{-4\mu}}{4}(1 + 3\bar{q})$, where $\bar{q}$ is the average similarity in the previous generation. If $\alpha$ and $\beta$ do not share a parent, which happens with probability *1-4/K*, then $< q_{\alpha\beta} >= e^{-4\mu}\ \bar{q}$. Therefore, at generation *t+1* we get

$$\bar{q}_{t+1} = \frac{4}{K}\frac{e^{-4\mu}}{4}(1 + 3\bar{q}_t) + (1 - \frac{4}{K})e^{-4\mu}\ \bar{q}_t = e^{-4\mu}[(1 - \frac{1}{K})\bar{q}_t + \frac{1}{K}]\ . \qquad (4)$$

Setting $\bar{q}_{t+1} = \bar{q}_t = q_0$, we find the equilibrium at

$$q_0 = \frac{1}{1+4\mu K}. \qquad (5)$$

The approximation holds for $\mu$ and *1/K* much smaller than one, which is always the case for real populations. For example, with *K* = 250 and $\mu$ = 0.025, population similarity will reach an equilibrium with $q_0$= 0.038. If $q_{min} \leq q_0$, the genetic distribution reaches this equilibrium, and no species will arise. On the other hand, if the minimal similarity $q_{min}$satisfies $q_{min} > q_0$, sympatric speciation will occur and we can estimate the time for it to occur. Subtracting $\bar{q}_t$ from both sides of Equation (4) and approximating $\bar{q}_{t+1} - \bar{q}_t = \frac{dq}{dt}$, we get the differential equation

$$\frac{dq}{dt} = \frac{1}{K}(1 - \frac{q_t}{q_0}) \qquad (6)$$

with solution given by

$$q(t) \ = \ q_0 + (1 - q_0)\ e^{-t/Mq_0}. \qquad (7)$$

The time $\tau$ to speciation can be estimated as the time $q(t)$ takes to reach $q_{min}$. Setting $q(\tau) = q_{min}$we find

$$\tau = Mq_0 log(\frac{1-q_0}{q_{min}-q_0}). \qquad (8)$$

*Manzo-Pelliti model of allopatric speciation and definition of* $\tau_a$

Manzo and Pelliti (1994) extended the Derrida-Higgs model to describe populations geographically isolated in islands to investigate the possibility of allopatric speciation in the presence of gene flow. They considered two islands with *K* individuals each following the Derrida-Higgs model with sexual reproduction. The interaction between individuals of different islands occurs only via migration: after reproduction with partners of the same island, individuals can migrate to the other island with probability $\varepsilon$. The genetic variation of the populations is measured by the quantity $q_{\alpha\beta}$, which is the similarity between individuals $\alpha$ and $\beta$ on the same island, and a new quantity $p_{\alpha\beta}$ is introduced to measure the similarity of individuals belonging to different islands. Similarly to the Derrida-Higgs model, the description of dynamical properties of the system, such as the values of the similarities in the equilibrium, $q_0$ and $p_0$, can be obtained in the approximation of infinitely large genomes and in the regime of small mutation rate and large population (Manzo and Pelliti, 1994).

In special, Manzo and Pelliti (1994) were interested in exploring the feasibility of the regime where $p_0 < q_{min} < q_0$: there is no speciation in the islands and each one beares a single species, but the islands differentiate from each other, i.e., the species are endemic. Therefore, the differentiation occurs only due to the exchange of individuals between the islands. In our model (Fig. 1 in the main text), we are similarly interested in studying the speciation of the parasite purely induced by host-switching (equivalent to migration events in the two islands model). Then, we also adopt $q_{min} < q_0$to inhibit the speciation of parasites that would occur within hosts in the absence of host-switching (we use $q_{min} = 0.5q_0$). However, our model can not be further described by the expressions of Manzo-Pelliti because the host-switching probability depends on time and multiple hosts (islands) emerge over the simulation. Nevertheless, we still can use this description for the scenario where there is no migration/host-switching, i.e., the allopatric speciation. As follows, we demonstrate how to calculate the time for allopatric speciation, $\tau_a$, which is a parameter to rescale the input phylogenies in our simulations.

*Dynamics in the two-islands model*

In the case of two islands, we need to distinguish the similarities between individuals inhabiting the same island, $q$, or different islands, $p$, as stated previously. For individuals $\alpha$ and $\beta$ that belong to the same island, the average similarity $\bar{q}_{t+1}$ follows equation (4), $\bar{q}_{t+1} = e^{-4\mu}[(1-\frac{1}{K})\bar{q}_t + \frac{1}{K}] = e^{-4\mu}\, Q(\bar{q}_t, K)$. Otherwise, if they were born in different islands, their similarity evolves as $\bar{p}_{t+1} = e^{-4\mu}\bar{p}_t$since they do not have any parents in common. After reproduction, pair of individuals of the same island keep their original geographic relation if they do not migrate or both migrate, with probability $(1-\epsilon)^2 + \epsilon^2 \equiv a(\epsilon)$. The probability of a pair of individuals changing their geographic relative position is given by $2\epsilon(1-\epsilon) \equiv b(\epsilon)$, that accounts for one individual staying at the island and the other migrating (note that $a(\epsilon) + b(\epsilon) = 1$). Therefore, the dynamics of $q$ and $p$ is given by

$$q_{t+1} = a(\epsilon)\, e^{-4\mu}\, Q(q_t, K) + b(\epsilon)\, e^{-4\mu}\, p_t \qquad (9)$$

$$p_{t+1} = b(\epsilon)\, e^{-4\mu}\, Q(q_t, K) + a(\epsilon)\, e^{-4\mu}\, p_t \qquad (10)$$

where we omitted the bars for simplicity. For $\epsilon$, $\mu$ and 1/K all much smaller than 1, the equations can be approximated by

$$q_{t+1} = (1 - 2\epsilon - 4\mu - 1/K)q_t + 2\epsilon p_t \qquad (11)$$

$$p_{t+1} = 2\epsilon q_t + (1 - 2\epsilon - 4\mu)\ p_t. \qquad (12)$$

We obtain the coupled dynamical equations for *q(t)* and *p(t)* from equations 11 and 12 using the approximations $q_{t+1} - q_t = dq/dt$ and $p_{t+1} - p_t = dp/dt$, as done previously:

$$\frac{dq}{dt} = -(2\epsilon + 4\mu + 1/K)q(t) + 2\epsilon p(t) \qquad (13)$$

$$\frac{dp}{dt} = 2\epsilon q(t) - (2\epsilon + 4\mu)p(t). \qquad (14)$$

When in strict allopatry, $\epsilon = 0$, the similarity between islands is simply given by $p(t) = e^{-4\mu t}$, i.e., it depends only on the mutation rate. The time for the diversification due to geographical isolation $\tau_a$ can be calculated by $p(\tau_a) = e^{-4\mu\tau_a} = q_{min}$. The time for allopatry is then

$$\tau_a = \frac{1}{4\mu} log(\frac{1}{q_{min}}). \qquad (15)$$

The time for allopatry decreases with $q_{min}$: making the reproduction more restricted (increasing $q_{min}$) facilitates the differentiation between islands, which occurs in a smaller time.

In our simulations, we find the relation between the length of the branches in the host phylogeny and the time in generations by assuming that the smaller branch corresponds to the minimal time to parasites speciate due to geographical isolation.$\tau_a$ is calculated by equation (15) with the input parameters and $q_{min} = 0.5\ q_0$, with $q_0$ given by equation (5). Also, we consider that the first host species (the root of the phylogeny) must evolve for $\tau_a$ generations for the parasite populations to accumulate genetic diversity before the first splitting event.

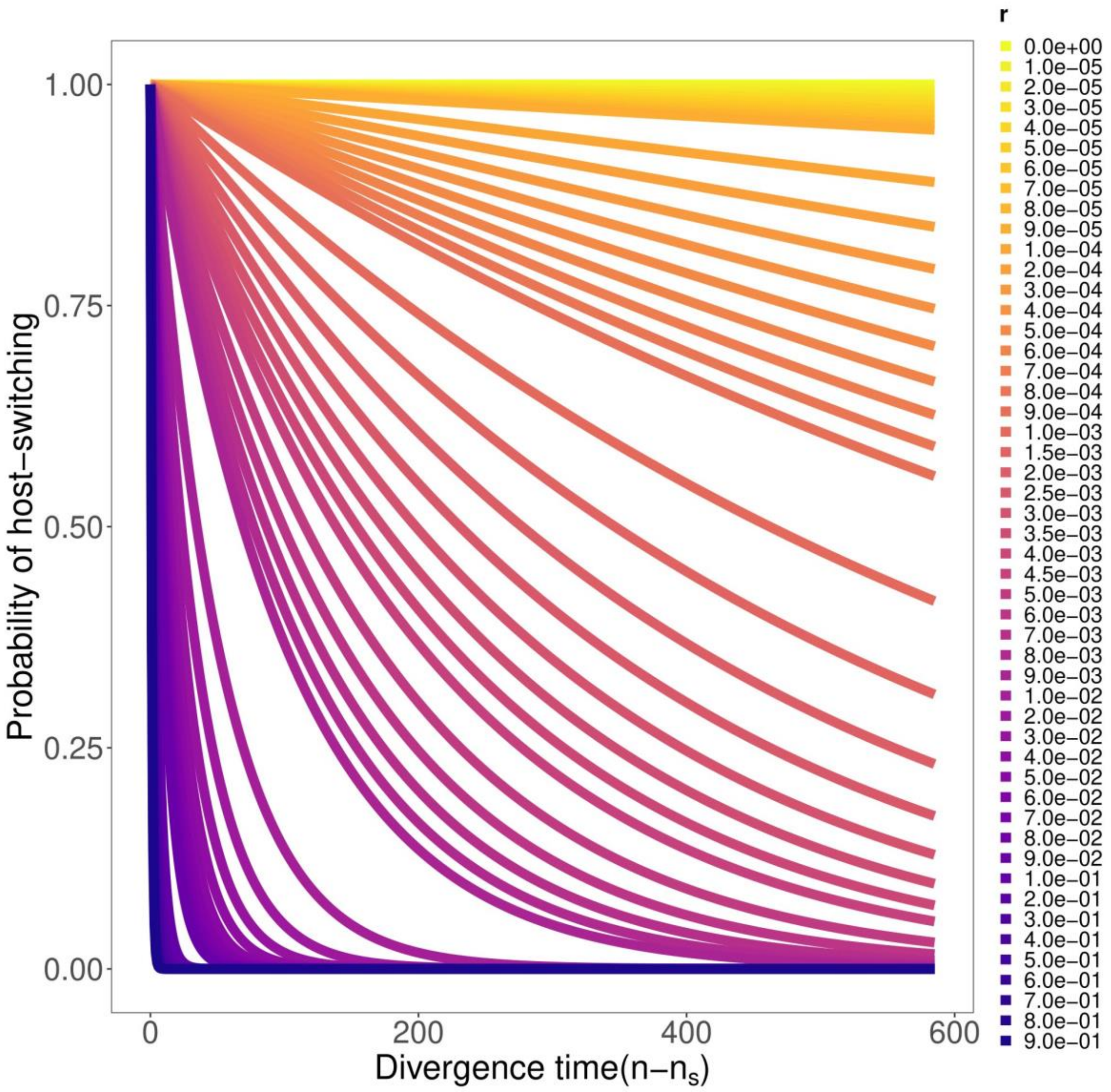


**Figure S1**. Probability of a parasite individual successfully host-switching from a host to another host species as a function of the divergence time (how long the two host species had diverged). If $r$=0 the probability is equal to 1 regardless of the divergence time, however this probability decreases if higher values of $r$ are considered.

*Parameters used*

The parameters involved in the model are described in Table S1. We considered four different carrying capacities $K = \{50,250,500,1000\}$. When we vary the population size (Figure S2 and S3) we do not observe differences in the patterns of beta diversity (β) and tree imbalance (normalised Sackin index - $I_n$) between the host-switching intensity, therefore we do not include the variation in the parasite species richness in our main results. Due to computational cost limitations, we selected the least rich host case (ID. 2) to show the influence of all population sizes and we had no qualitative difference between them (see species richness in Figure S4). The model is not too sensitive in response to changes in parasites' population size, which allows us to fix the values. Then, for the parasite population, we selected $K = 250$. Similarly, we also choose the $\mu = 0.025$, to carry out the simulations with different scenarios of host-switching intensity. We replicated each combination of the parameters for each empirical case 50 times. We included an initial transition time using the value of allopatric time in the model to eliminate the effect of the initial condition on evolutionary patterns. $t$ represent the evolutionary time of the parasites was parameterized according to the evolutionary time of the hosts and the time of allopatric speciation for each empirical case. Simulations were performed separately for each host phylogenies (Table S1).

**Table S1**. Model parameters with a short description and the investigated values.

| Parameters | Short definition | Investigated values |
|---|---|---|
| $K$ | Carrying capacity of parasite species per host species. | 50, **250**, 500, 1000 |
| $\mu$ | Mutation rate. | 0.001, **0.025** |
| $q_{min}$ | Minimal genetic similarity. | **0.5*$q_0$** |
| $r$ | Intensity of decline in host-switching probabilities as the phylogenetic distance increases between hosts. | **0-1** |
| | Number of simulation repetitions for a given set of parameters. | **50** |

| | |
|---|---|
| Total number of iterations | ID 1. 1812*<br>ID 2. 1491*<br>ID 3. 1343*<br>ID 4. 702*<br>ID 5. 624*<br>ID 6. 541*<br>ID 7. 2614*<br>ID 8. 339*<br>ID 9. 248* |

Note: The bold values are the fixed values used in the presented results, while the other parameters used for the sensibility test.*total time for each empirical case (community).

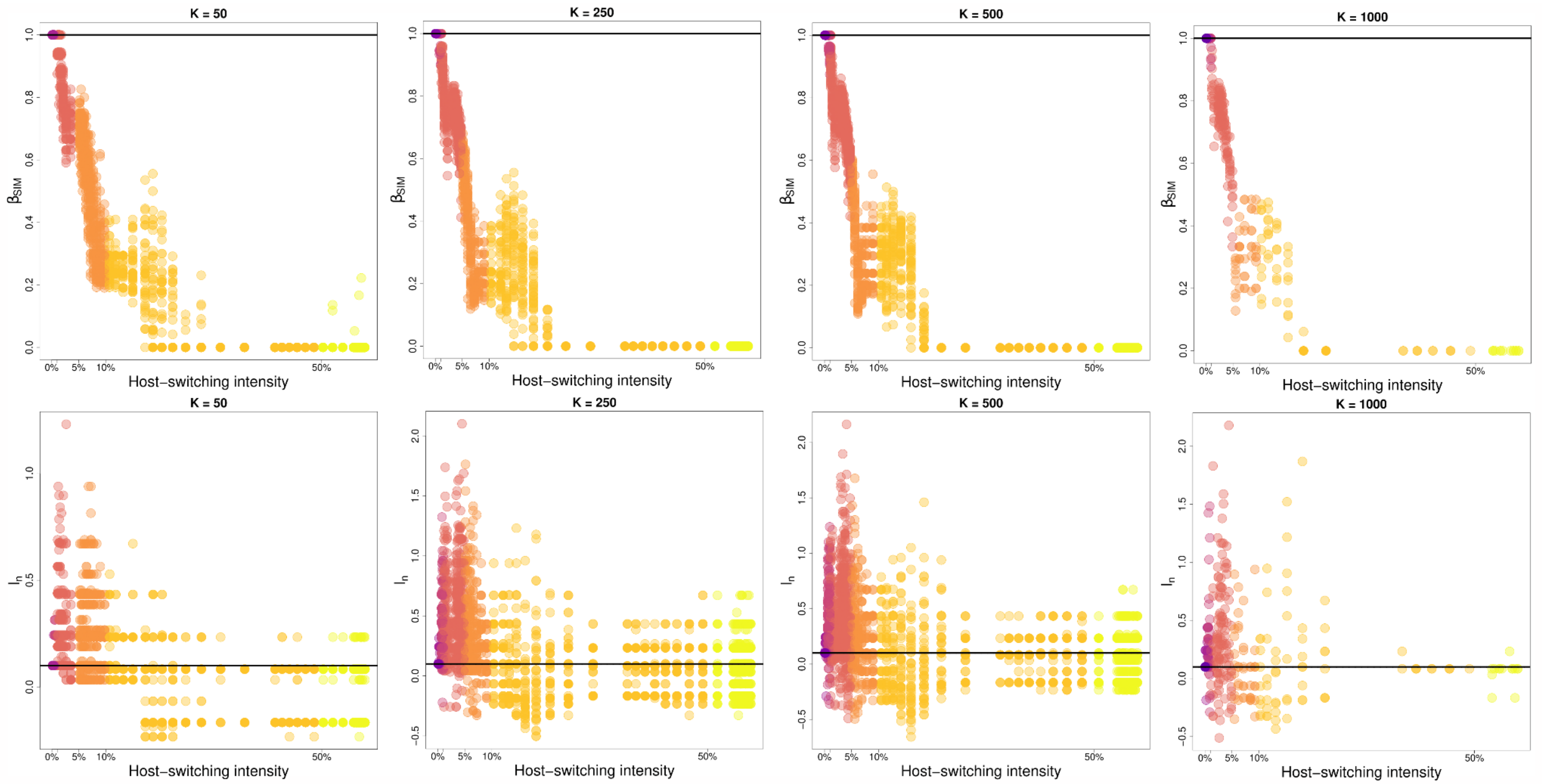


**Figure S2**. Population size test. Relationship between variation in the composition and host-switching intensity (a-d) and the tree imbalance (normalised Sackin index - $I_n$) and host-switching intensity (e-h) for different population sizes for empirical case ID. 2. Each *K* represents the population size tested. The lines refer to empirical information of parasite (continuous) and host (dotted). For this case, these lines overlap. The colored dots are redundant with the x-axis scale of graphs, but are intended to guide the interpretation. For each configuration of the parameters of host-switching intensity a total of 50 runs were performed for each carrying capacity, with the exception of K = 1000, which demanded more time for running. For this case, a total of 10 runs were performed.

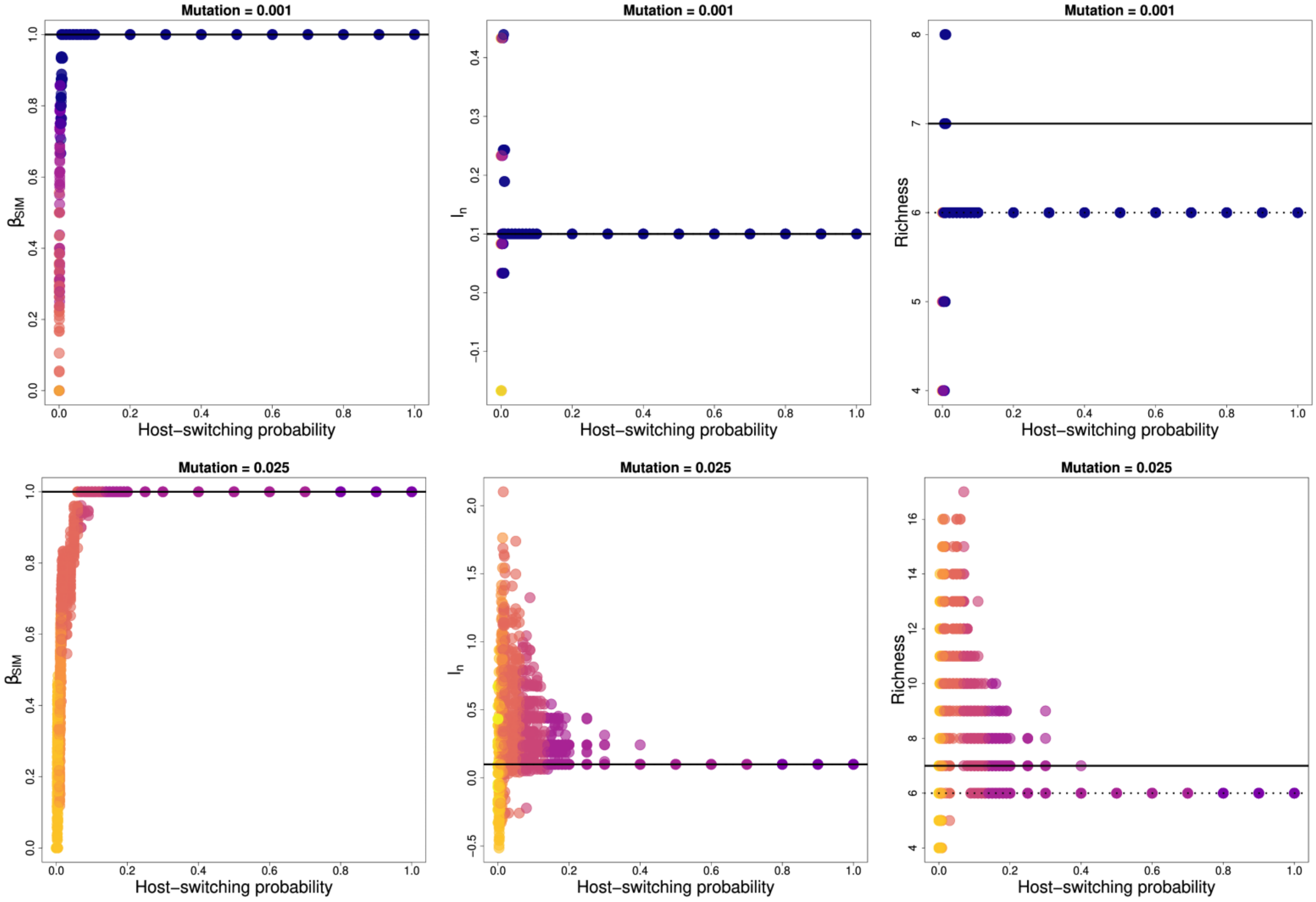
Mutation = 0.001
Mutation = 0.001
Mutation = 0.001
Mutation = 0.025
Mutation = 0.025
Mutation = 0.025
βSIM
In
Richness
Host−switching probability
0.0
0.2
0.4
0.6
0.8
1.0

**Figure S3**. Mutation test. Relationship between variation in the composition and host-switching intensity (a-d) and the tree imbalance (normalised Sackin index - $I_n$) and host-switching intensity (e-h) for two mutation rates for empirical case ID. 2. Each $\mu$ represents the mutation rate tested. The lines refer to empirical information of the parasite (continuous) and host (dotted). For this case, these lines overlap. The colored dots are redundant with the x-axis scale of graphs but are intended to guide the interpretation. For each configuration of the parameters of host-switching intensity a total of 50 runs were performed for each carrying capacity, with K = 250.

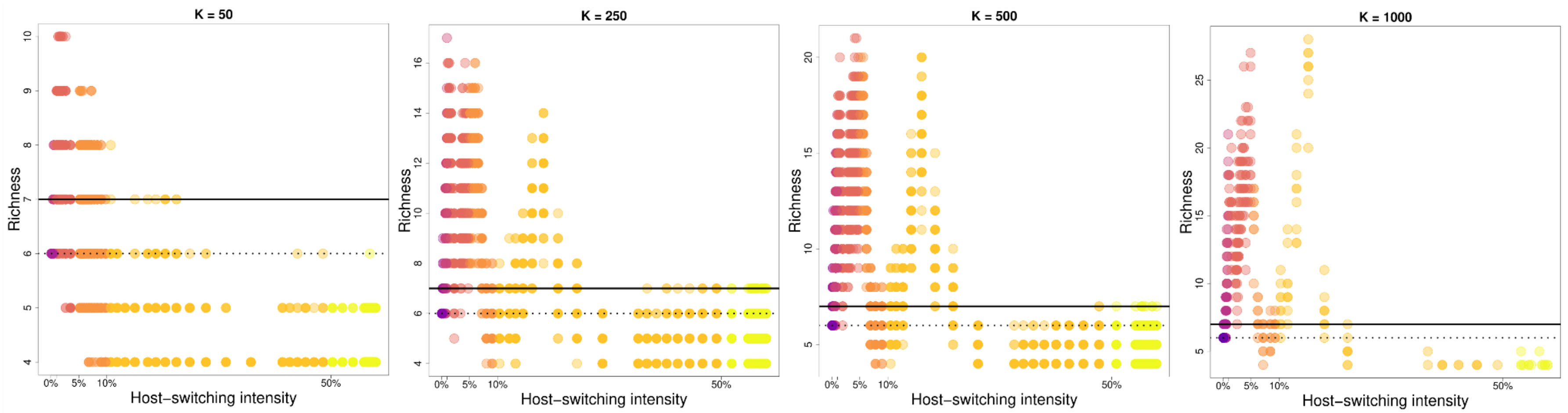


**Figure S4**. Relationship between species richness and host-switching intensity (a-d) for empirical case ID. 2. Each *K* represents the population size tested. The lines refer to empirical information of parasite (continuous) and host (dotted). The colored dots are redundant with the x-axis scale of graphs, but are intended to guide the interpretation. For each configuration of the parameters of host-switching intensity a total of 50 runs were performed for each carrying capacity, with the exception of K = 1000, which demanded more time for running. For this case, a total of 10 runs were performed.

*Details for empirical database*

To test our model, the cases (communities) essentially need to have real phylogenies for hosts and parasites and their interactions. A literature search was performed using the phrase "*phylogeny host parasite*" was carried out using Google Scholar between December 1, 2019 and January 2021, which identified more than 10,000 works. Among these, a total of 100 articles were selected for feasibility analyses for testing the model. Articles that did not contain phylogeny were immediately excluded. Studies focussing on the population level were also excluded. Studies that included less than six taxa were excluded because, as they have a low sampling effort, they can lead to misinterpretations. Additionally, studies with species that have asexual reproduction were excluded from consideration, as our model is sexed. Finally, for inclusion in our analyses we extracted nine cases (Table 1 and Figure 2 in main manuscript and Figure S5-S13) with knowledge of the associations and phylogenies that exist between parasites and host tips. Then, we studied the effects of host-switching and host evolution signatures on parasite speciation patterns (phylogenetic trees and variation in species composition) only for surviving species. Extinct species were not included in the analyses. The interactions of all analysed cases are available at https://github.com/elviradbastiani/host_switching_model.

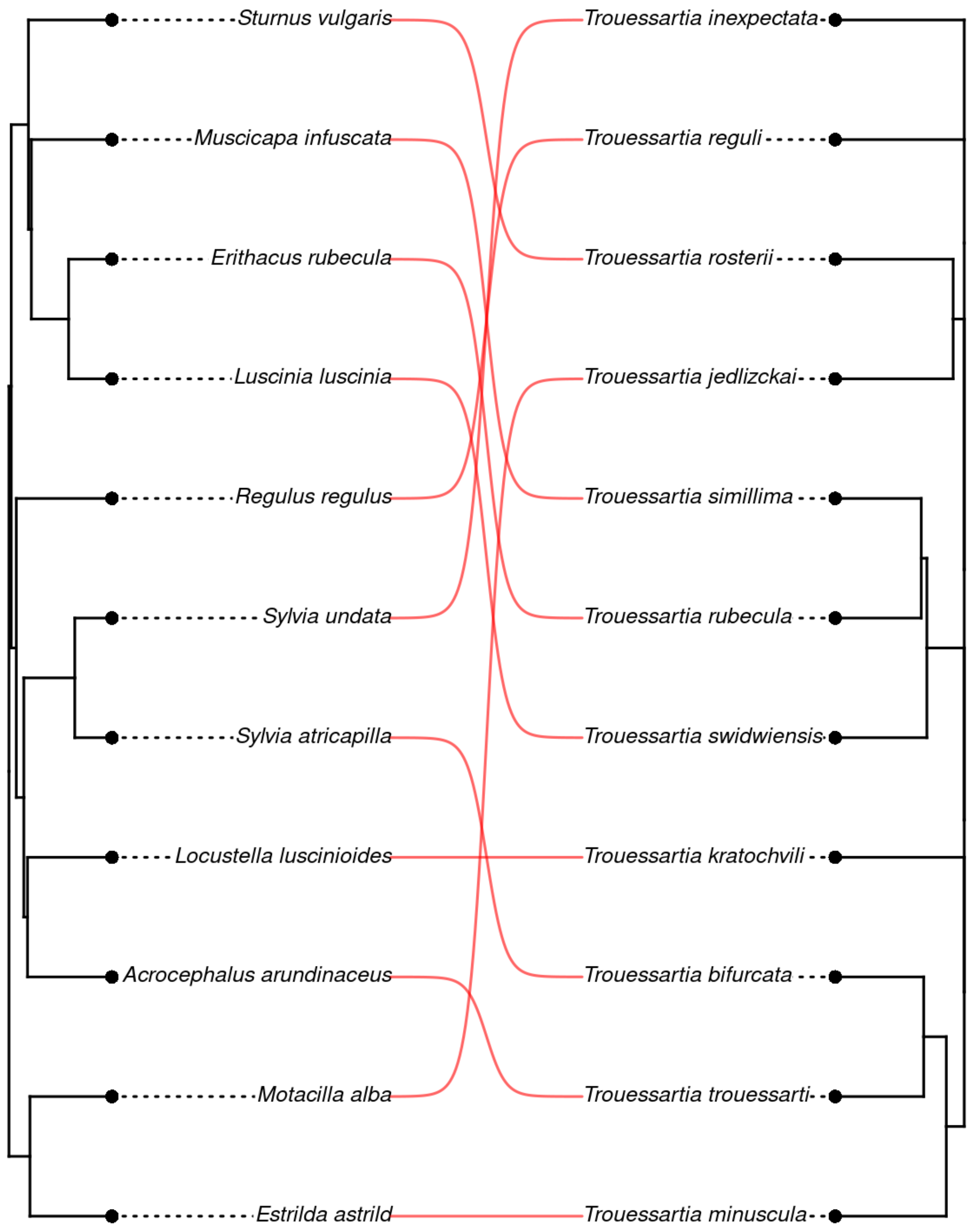


**Figure S5**. Phylogenetic trees that correspond to the empirical data of Bird and feather mites (*Trouessartia* spp.) corresponding to ID. 1 extracted from Donã et al. 2017.

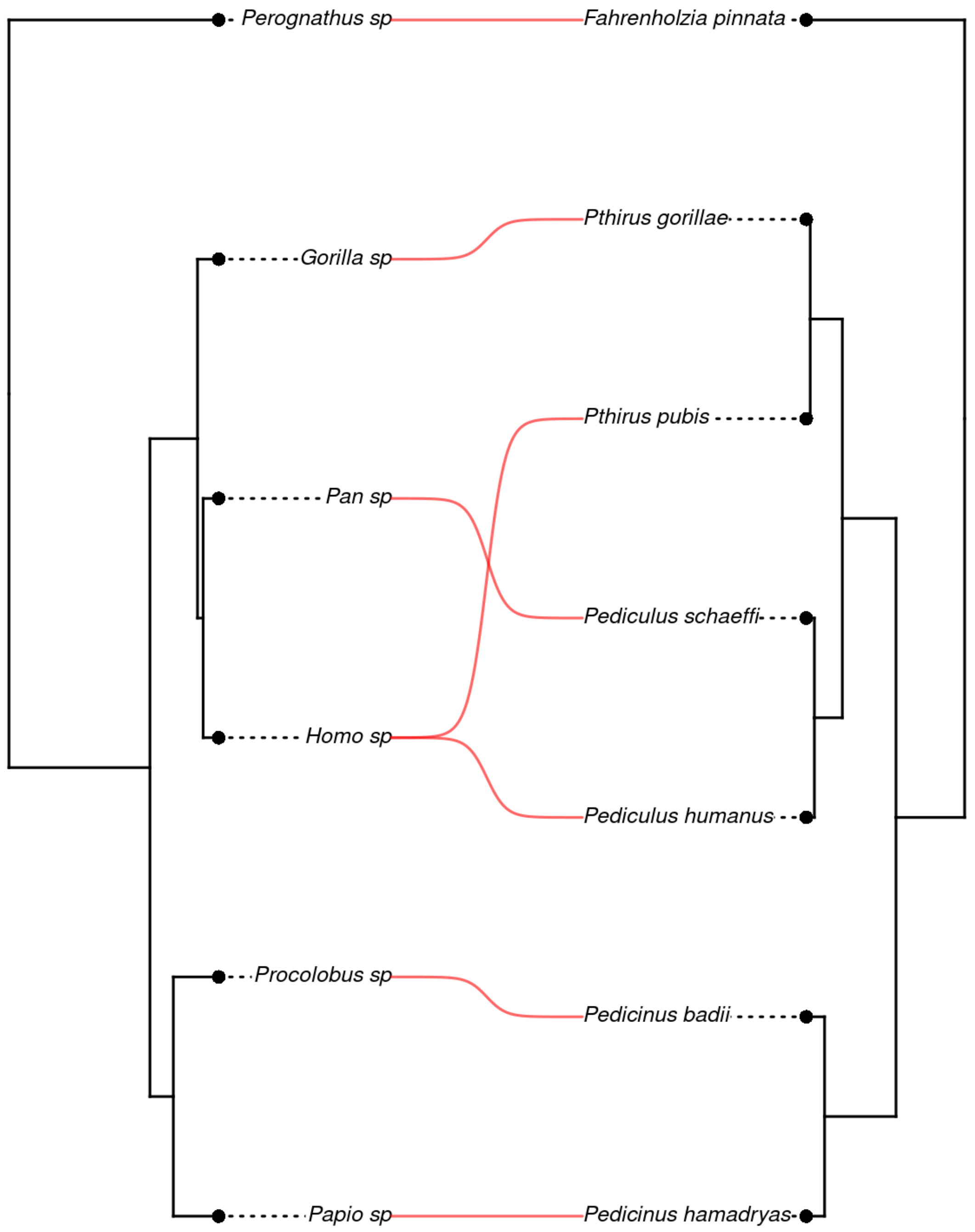


**Figure S6**. Phylogenetic trees that correspond to the empirical data of Mammals and Lice (*Pediculus* spp. and *Pthirus* spp.) corresponding to ID. 2 extracted from Reed et al. 2007.

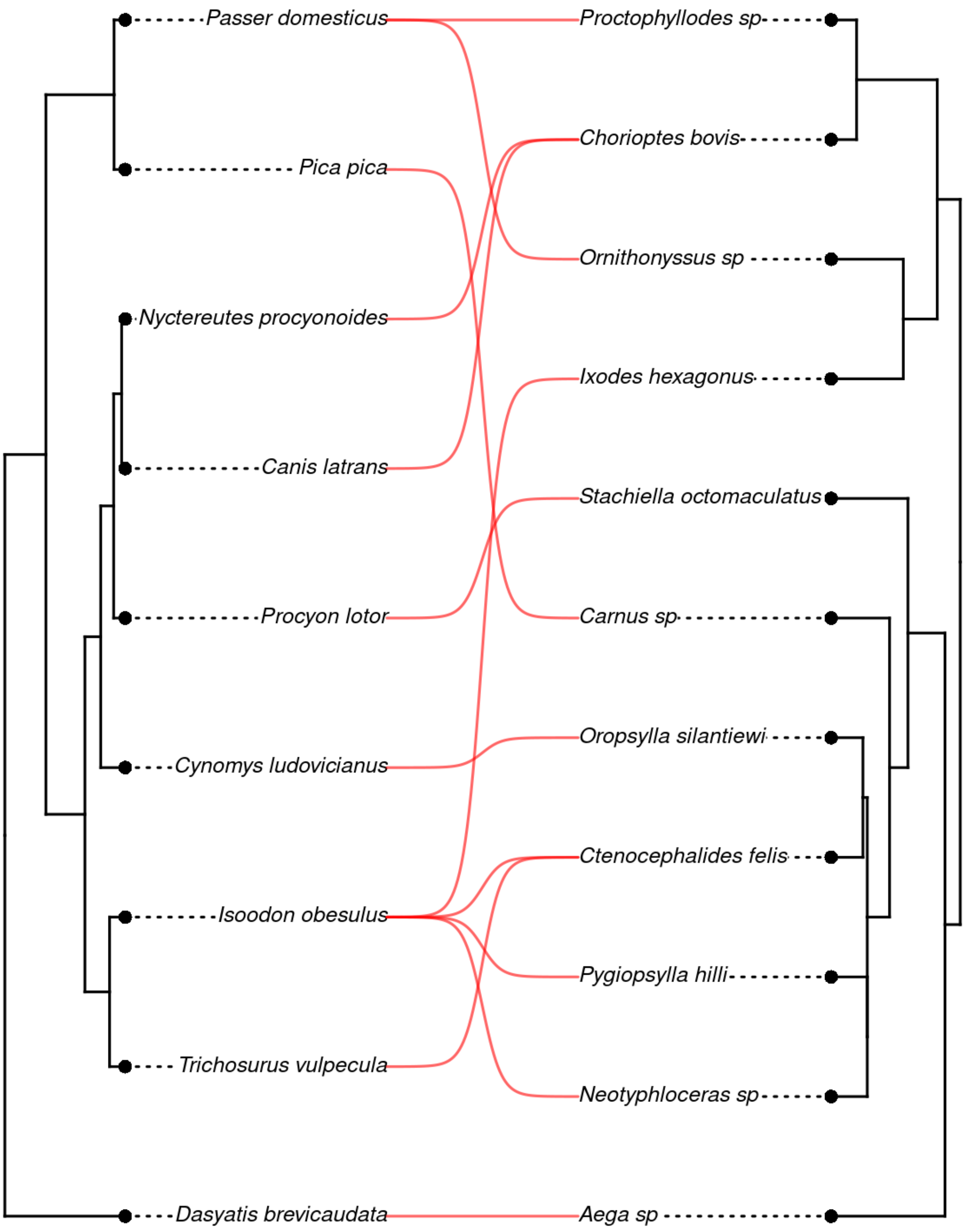


**Figure S7**. Phylogenetic trees that correspond to the empirical data of wildlife and arthropods corresponding to ID. 3 extracted from Becker et al. 2018.

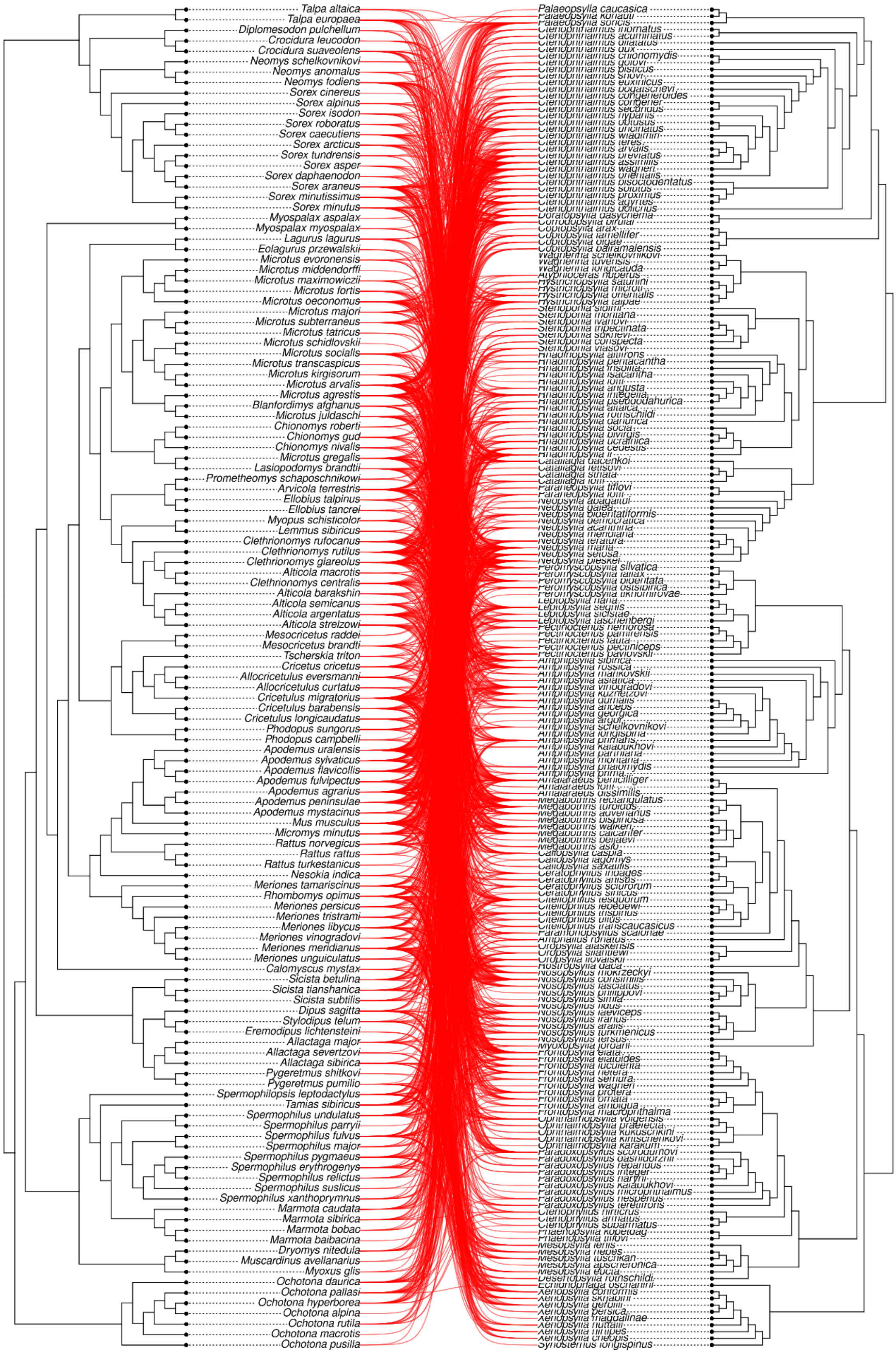

**Figure S8**. Phylogenetic trees that correspond to the empirical data of rodents and fleas corresponding to ID. 4 extracted from Krasnov et al. 2016.

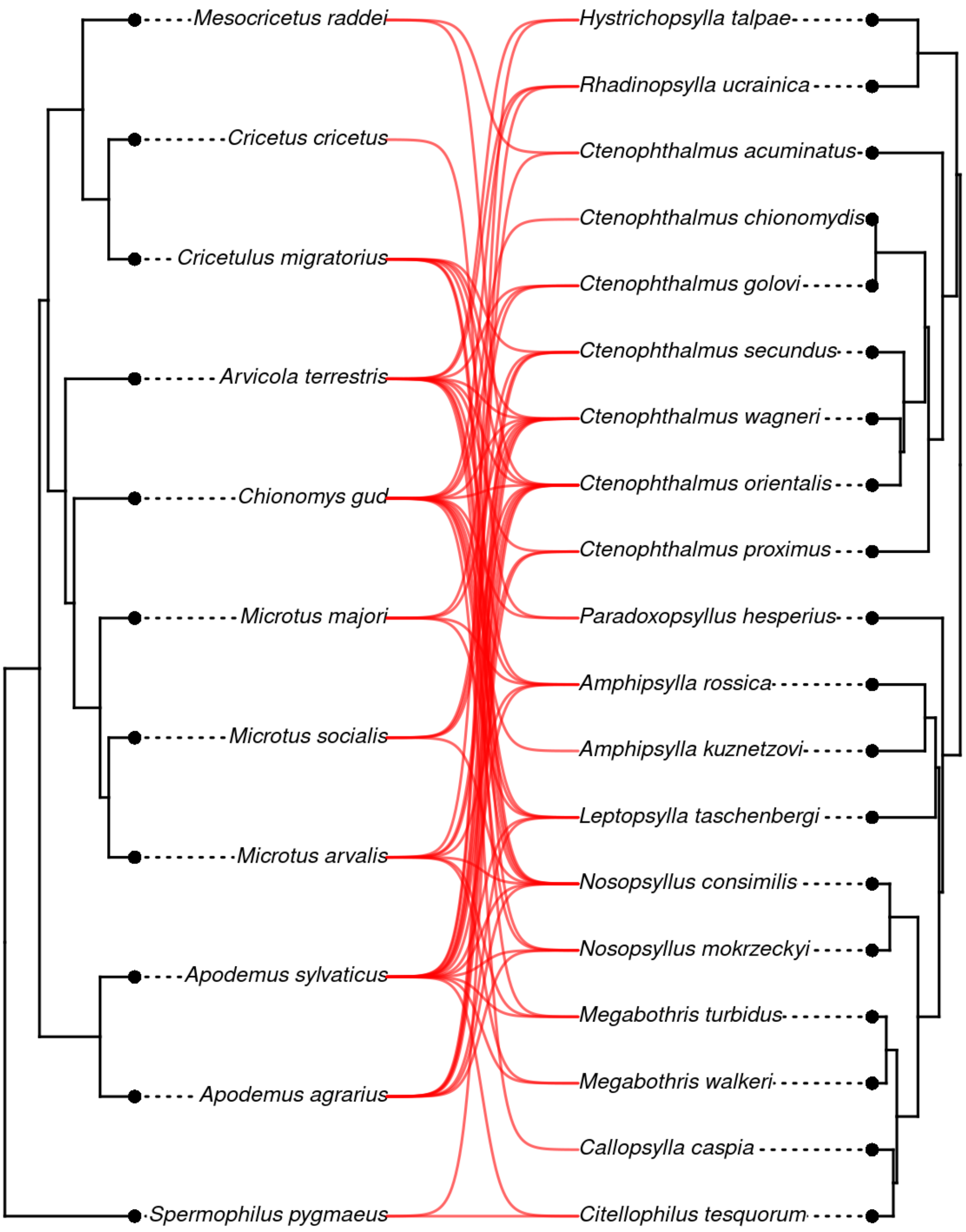


**Figure S9**. Phylogenetic trees that correspond to the empirical data of rodents and fleas corresponding to ID. 5 extracted from Krasnov et al. 2016.

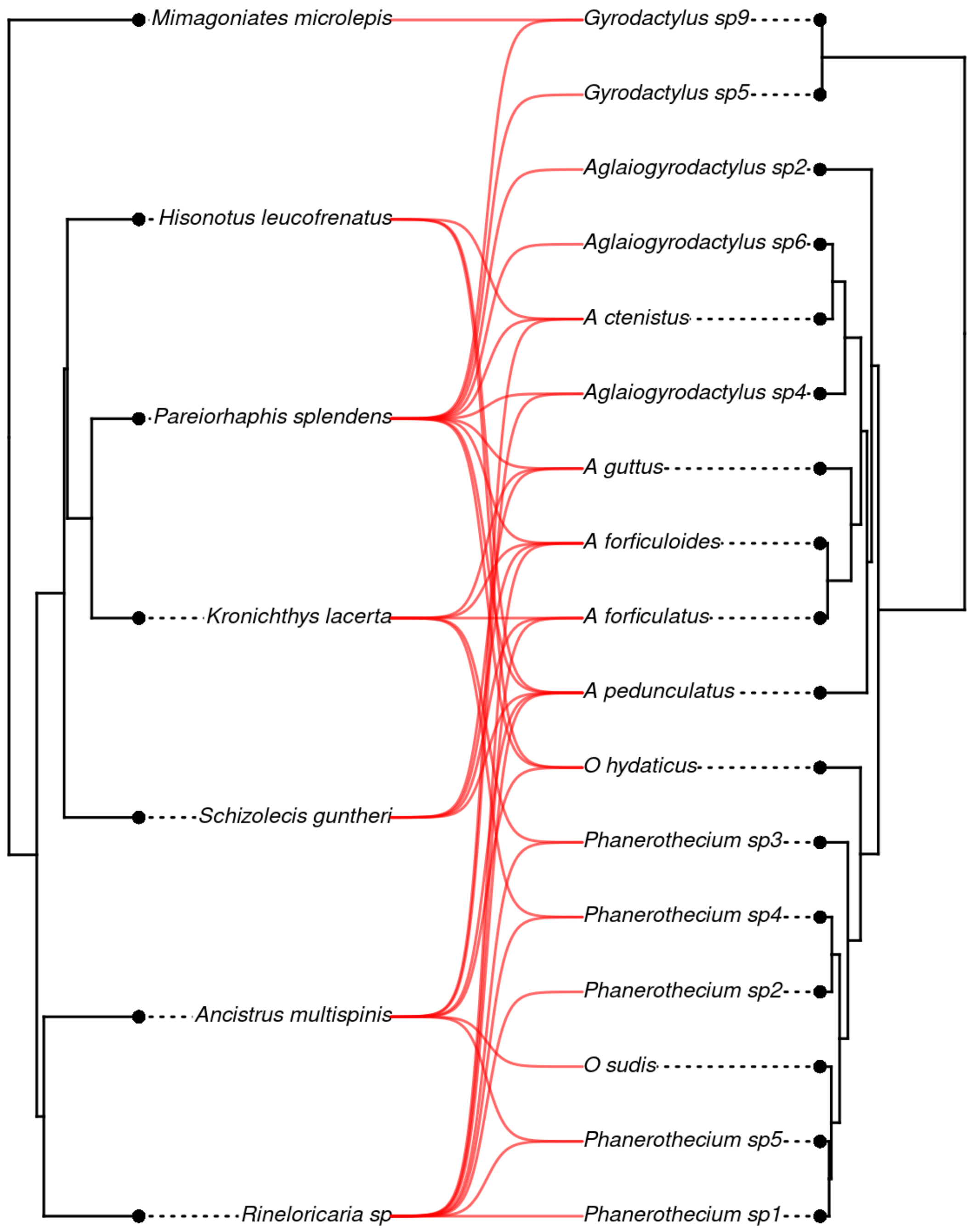


**Figure S10**. Phylogenetic trees that correspond to the empirical data of fish and platyhelminthes (Gyrodactylidae) corresponding to ID. 6 extracted from Patella et al. 2017.

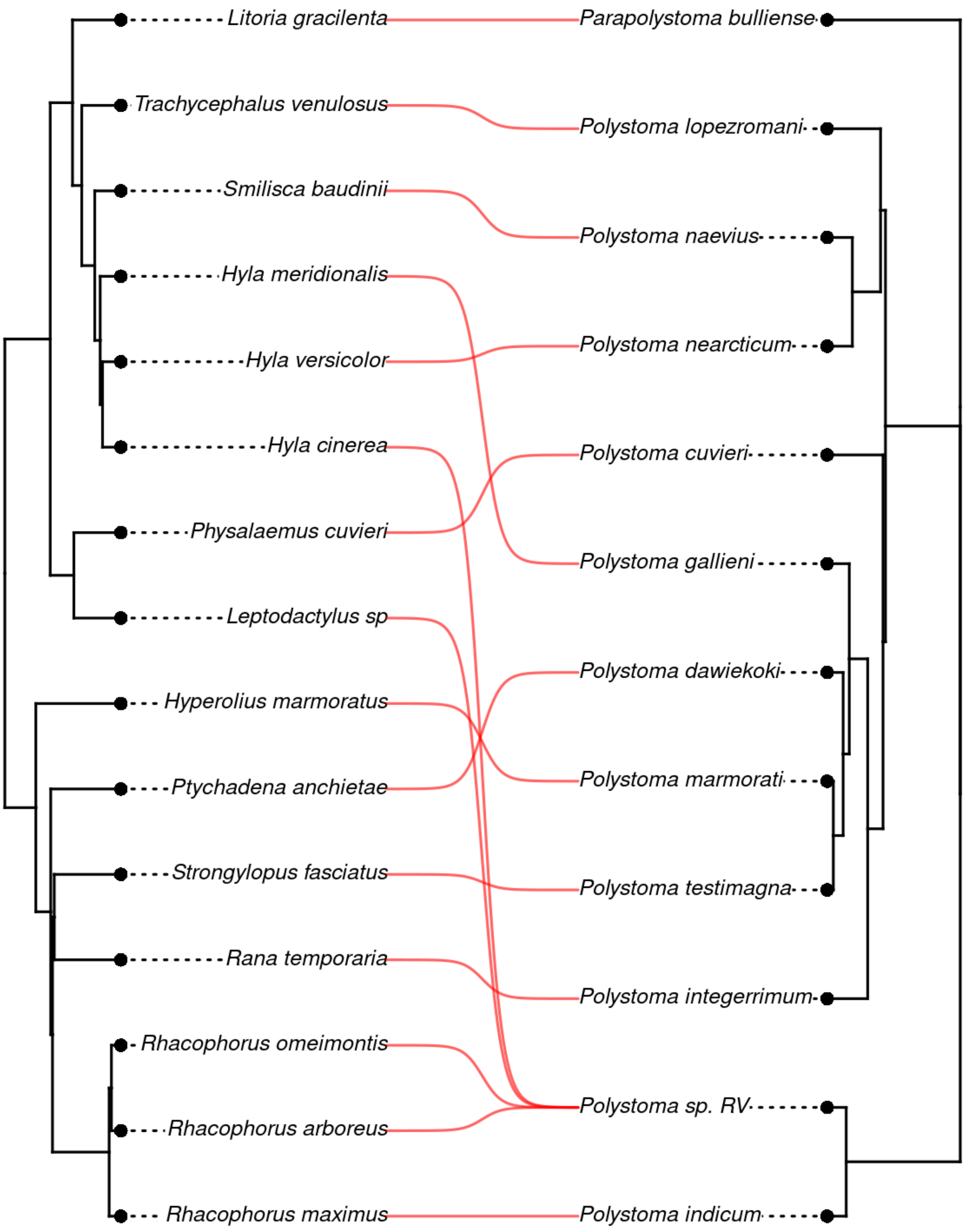


**Figure S11**. Phylogenetic trees that correspond to the empirical data of frogs and polystomes (Polystomatidae) corresponding to ID. 7 extracted from Badets et al. 2011.

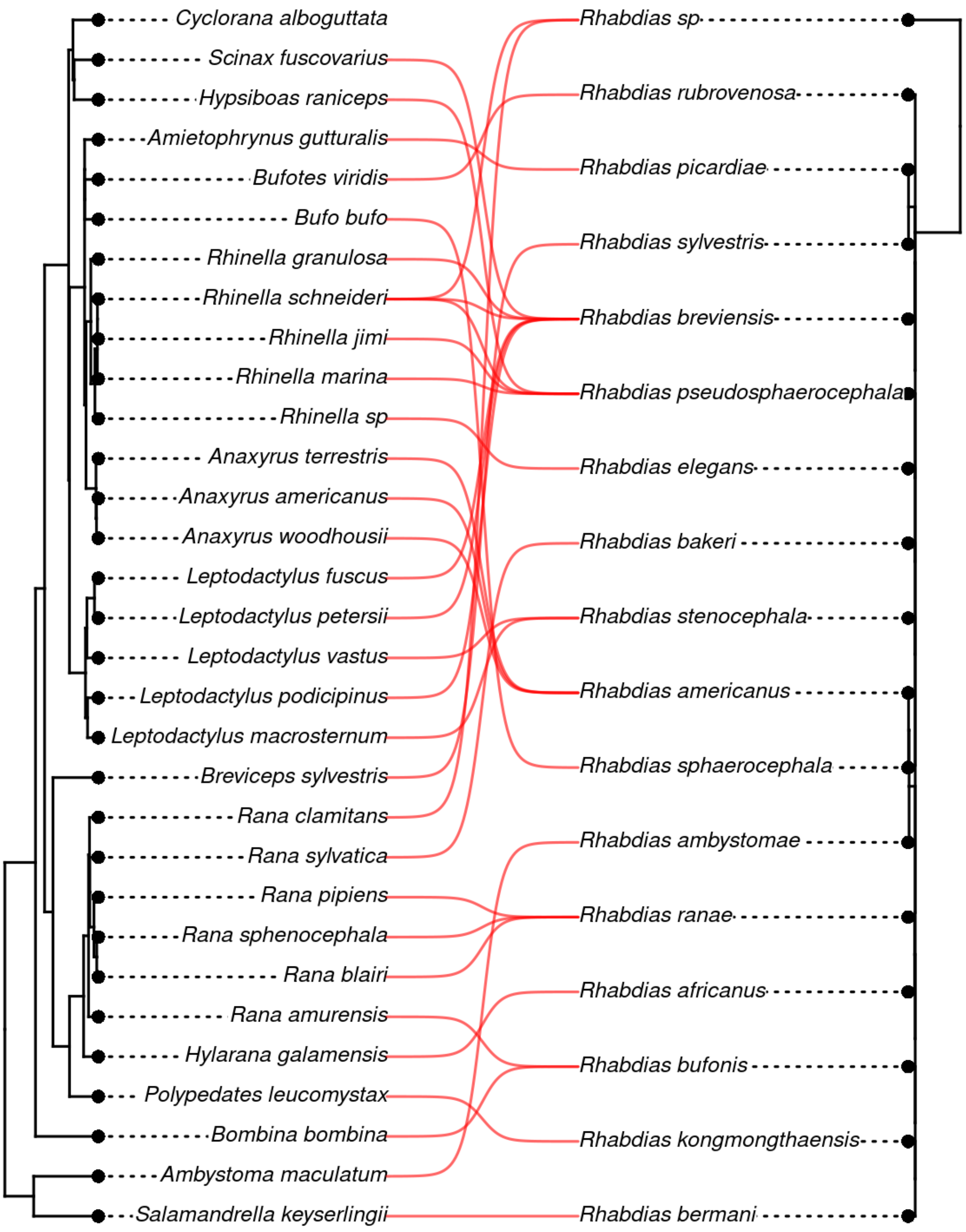


**Figure S12**. Phylogenetic trees that correspond to the empirical data of frogs and Nematodes (*Rhabdias* spp. ) corresponding to ID. 8 extracted from Müller et al. 2018.

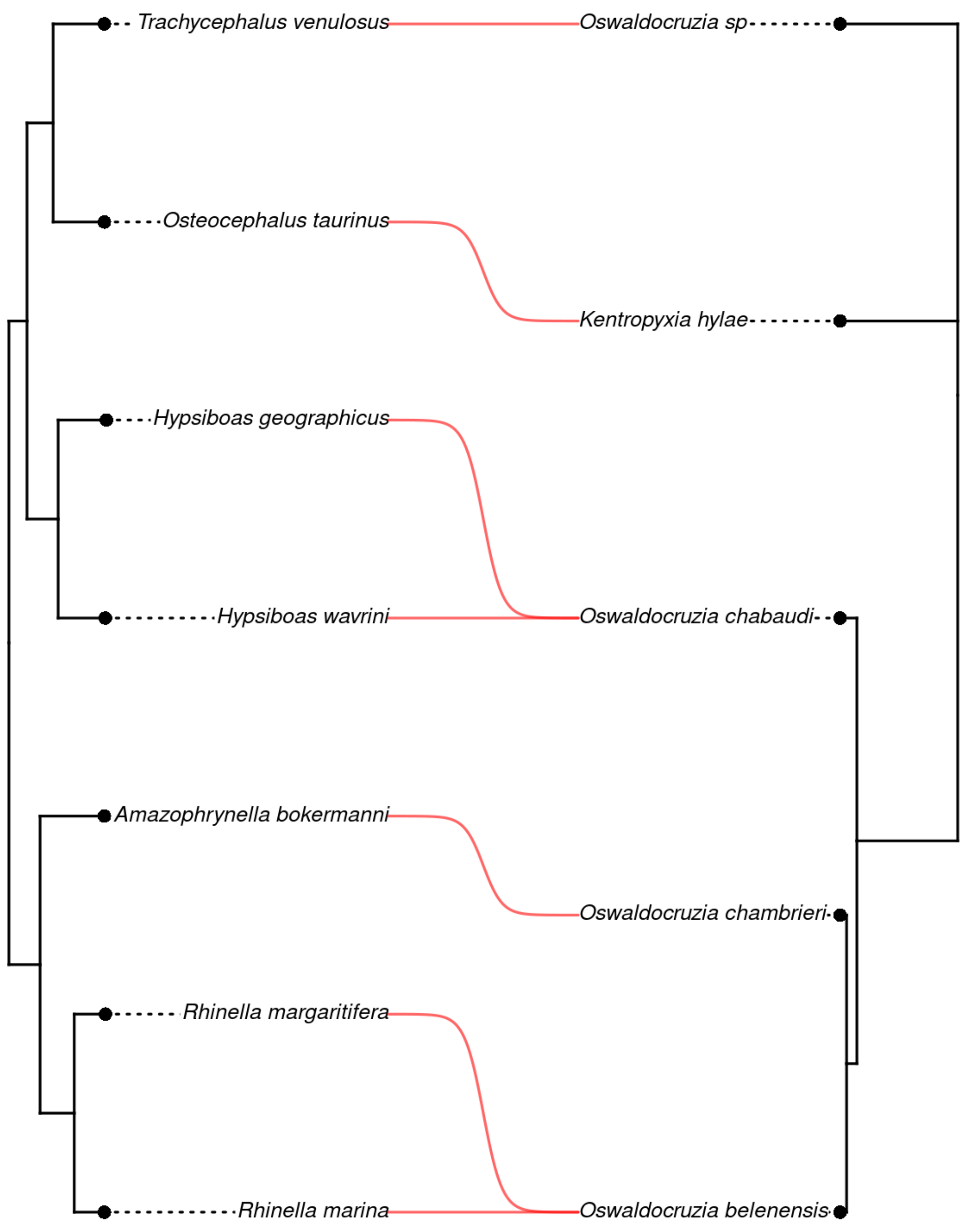


**Figure S13**. Phylogenetic trees that correspond to the empirical data of frogs and Nematodes (*Oswaldocruzia* spp. ) corresponding to ID. 8 extracted from Willkens et al. 2021.

*Specific results*

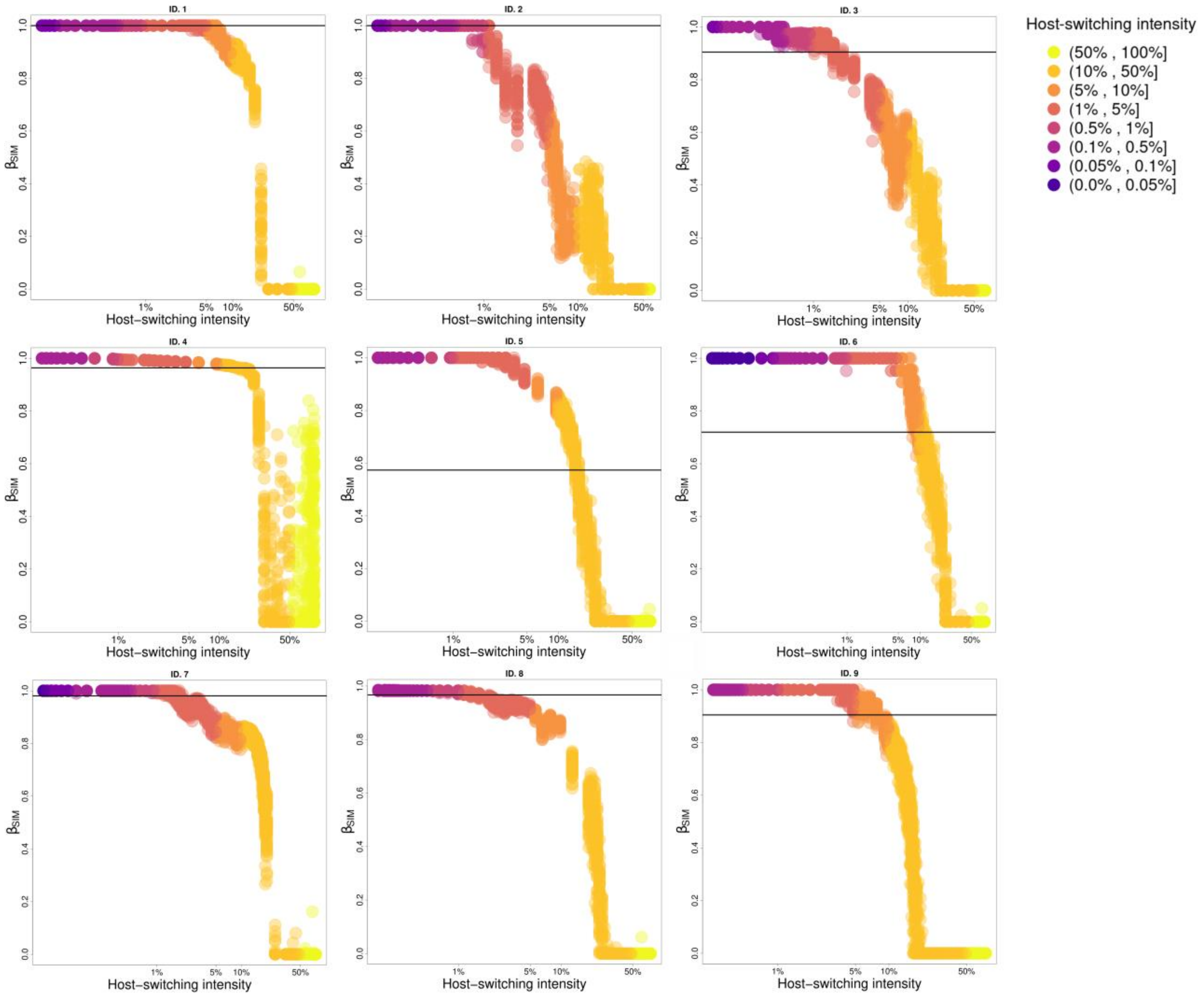
ID. 1
ID. 2
ID. 3
ID. 4
ID. 5
ID. 6
ID. 7
ID. 8
ID. 9
$\beta_{SIM}$
Host–switching intensity
1%
5%
10%
50%
Host-switching intensity
(50% , 100%]
(10% , 50%]
(5% , 10%]
(1% , 5%]
(0.5% , 1%]
(0.1% , 0.5%]
(0.05% , 0.1%]
(0.0% , 0.05%]

**Figure S14**. The relationship between variation in parasite composition (measured by the metric beta diversity - β - y-axis) and intensity of host-switching (x-axis and colours) for the nine empirical cases. A total of 50 runs were performed for each configuration of the parameter "host-switching intensity", with K=250 individuals. We present the results for endoparasites and ectoparasite empirical cases (continuous line refers to parasite information, and dotted line refers to hosts). To help distinguish the different intensity of the host switch, we make colour scales for each percentage interval.

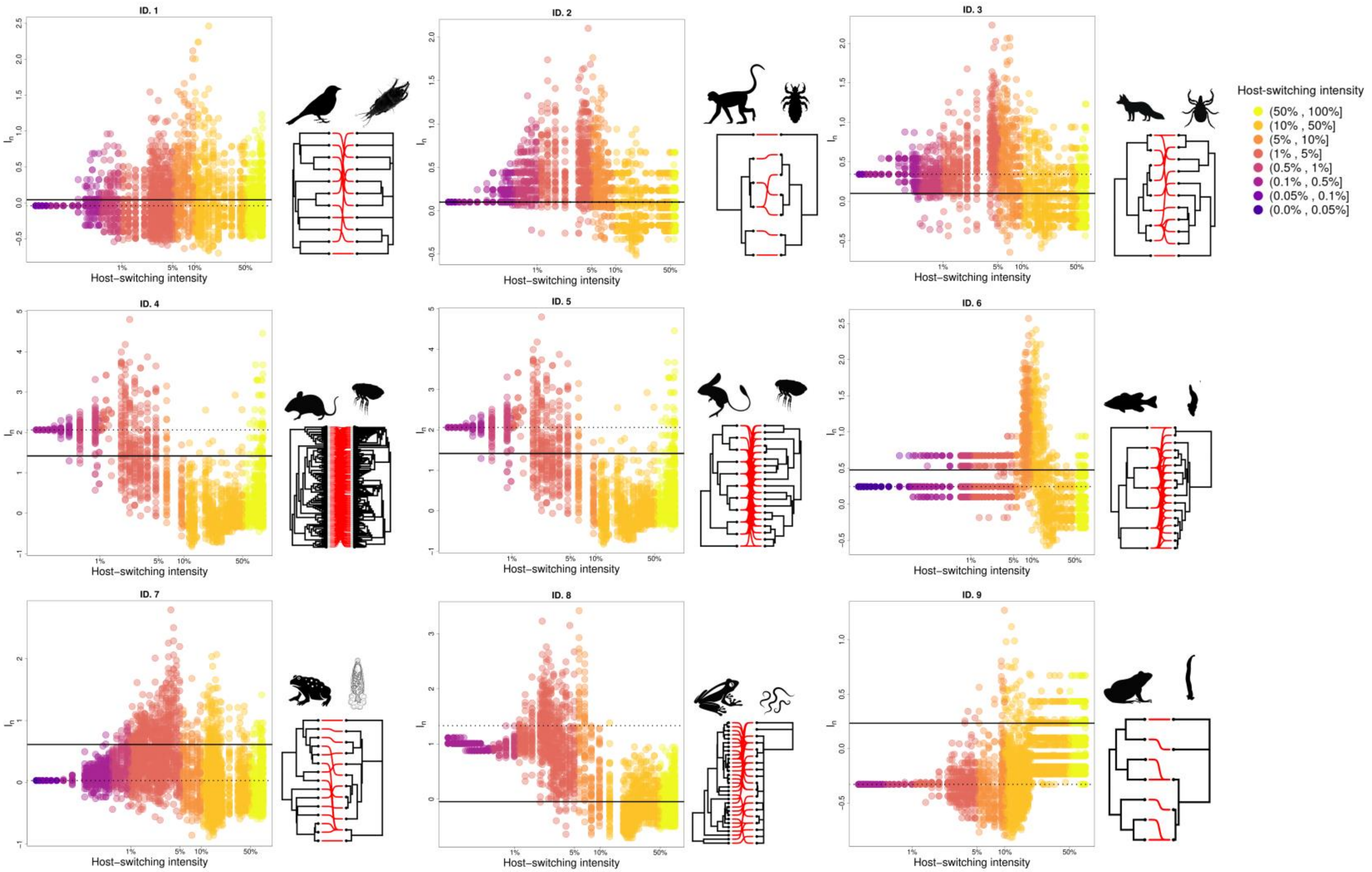
ID. 1
ID. 2
ID. 3
ID. 4
ID. 5
ID. 6
ID. 7
ID. 8
ID. 9
$I_n$
Host–switching intensity
1%
5%
10%
50%
Host-switching intensity
(50% , 100%]
(10% , 50%]
(5% , 10%]
(1% , 5%]
(0.5% , 1%]
(0.1% , 0.5%]
(0.05% , 0.1%]
(0.0% , 0.05%]

**Figure S15**. The relationship between variation in parasite composition (measured by the metric normalised Sackin index - $I_n$ - y-axis) and intensity of host-switching (x-axis and colours) for the nine empirical cases. A total of 50 runs were performed for each configuration of the parameter "host-switching intensity", with K=250. We present the results for endoparasites and ectoparasite empirical cases (continuous line refers to parasite information, and dotted line refers to hosts). To help distinguish the different intensities of the host switch, we make colour scales for each percentage interval.

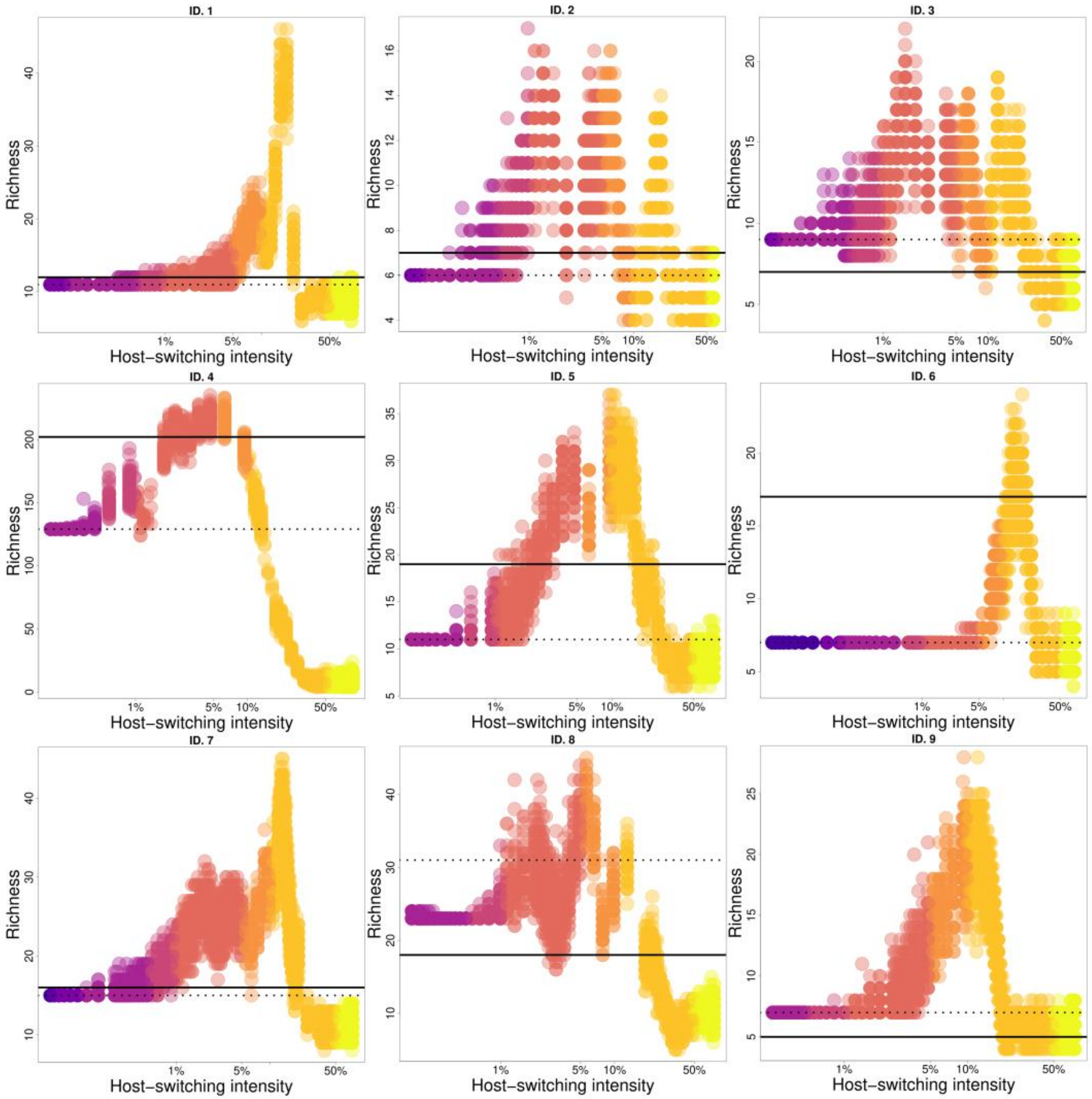
ID. 1
ID. 2
ID. 3
ID. 4
ID. 5
ID. 6
ID. 7
ID. 8
ID. 9
Richness
Host–switching intensity
1%
5%
10%
50%

**Figure S16**. The relationship between variation in parasite composition (measured by the metric normalised Sackin index - $I_n$ - y-axis) and intensity of host-switching (x-axis and colours) for the nine empirical cases. A total of 50 runs were performed for each configuration of the parameter "host-switching intensity", with K=250. We present the results for endoparasites and ectoparasite empirical cases (continuous line refers to parasite information, and dotted line refers to hosts). To help distinguish the different intensities of the host switch, we make colour scales for each percentage interval.

*Example eco-evolutionary patterns over time*

Here we illustrated the complete temporal results (Fig. S17 and S18) only the empirical case of the feather mites associated with birds to explain the ecological and evolutionary trajectories over time. In the empirical case represented in Fig. 1 in the main document (feather mites associated with birds - ID. 1), all host species arose only in evolutionary time 1500. Only after this time, resource limitations become more evident and interactions start to become more restricted. For this case, as hosts diversify, they arrive at an end time (1812) with an eco-evolutionary pattern similar to the interactions in the empirical study. In the other cases we observe similar patterns.

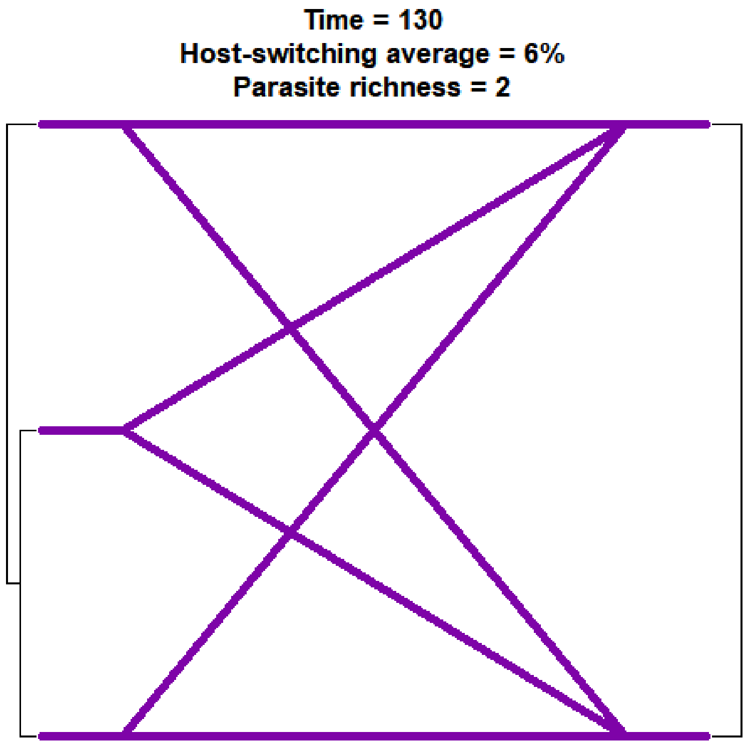


**Figure S17**. Complete temporal results (GIF) show the patterns of host-switching over evolutionary time for feather mites associated with birds.

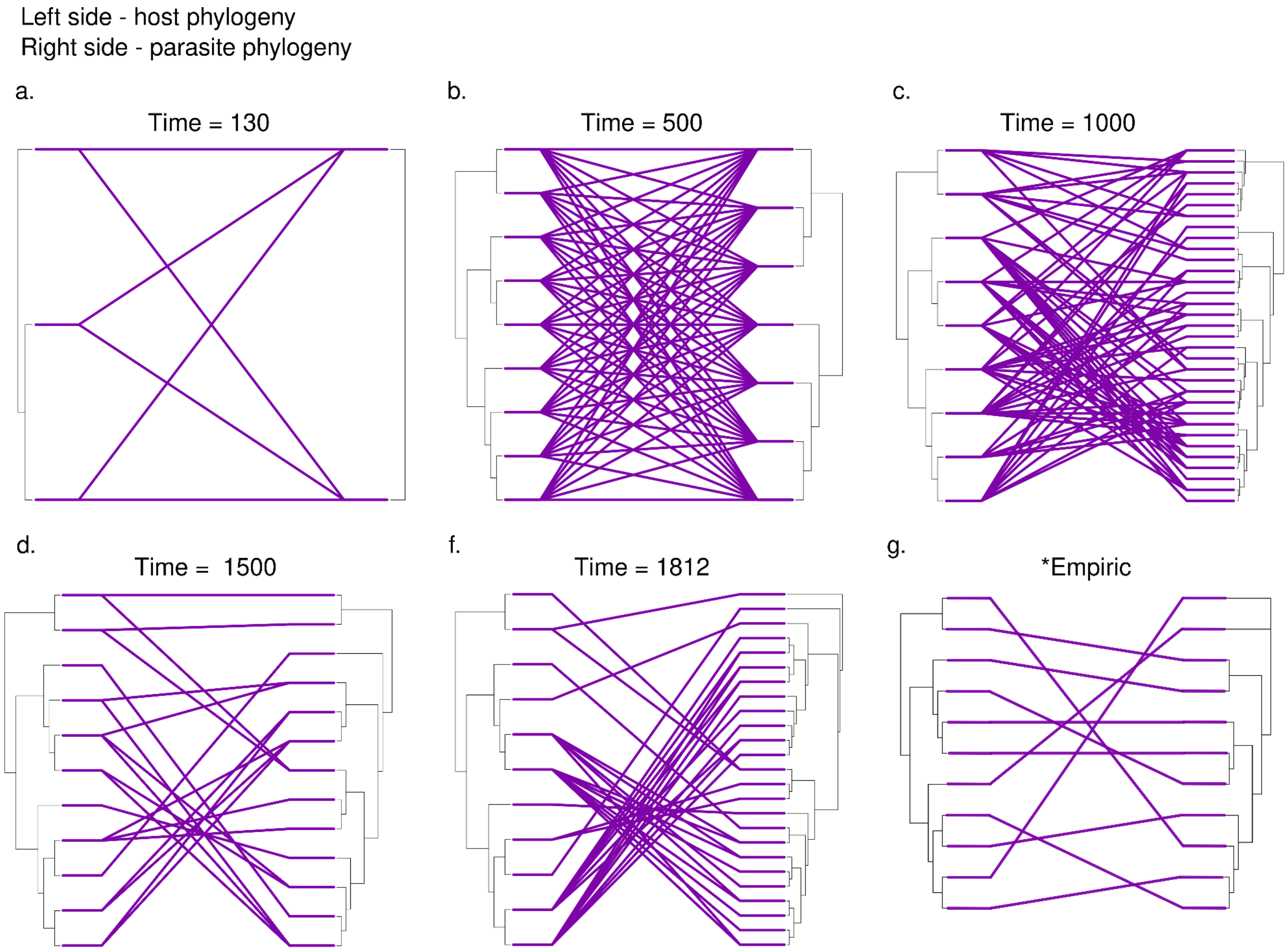
Left side - host phylogeny
Right side - parasite phylogeny
a.
Time = 130
b.
Time = 500
c.
Time = 1000
d.
Time = 1500
f.
Time = 1812
g.
*Empiric

**Figure S18**. Ecological and evolutionary patterns of host and parasite to empirical study (case ID1 - feather mites associated with birds) over time (i-vi) with host-switching intensity between 1% - 5%. The end time has similar structures with case study 1. *empirical phylogeny host-parasite.

[Dataset] Willkens Y., Furtado A. P., Dos Santos J. N.  de Vasconcelos Melo F. T. 2021. Do host habitat use and cospeciation matter in the evolution of *Oswaldocruzia* (Nematoda, Molineidae) from neotropical amphibians? Journal of Helminthology, 95, e33. Accession number: EDB_10.1017/S0022149X21000250
